# Scaling data analyses in cellular cryoET using comprehensive segmentation

**DOI:** 10.1101/2025.01.16.633326

**Authors:** Mart G. F. Last, Lenard M. Voortman, Thomas H. Sharp

## Abstract

Automation and improved hardware have greatly accelerated the rate of data generation in cryoET. As the field moves towards *quantitative cryoET*, the scale of the resulting datasets presents a significant challenge for analysis and interpretation. To explore ways of handling datasets comprising thousands of tomograms, we investigated a comprehensive segmentation strategy – assigning an ontology-based identity to every voxel in a dataset – that is based on the sequential application of multiple convolutional neural networks. Using an openly available dataset of over 1800 *Chlamydomonas reinhardtii* tomograms as a test case, we demonstrate the segmentation of 25 different subcellular features across the full dataset, while requiring only a few seconds of processing time per tomogram. We show how the approach enables the representation of large datasets as searchable databases and propose the usage of ontology-based segmentations for improving two common processing tasks in cryoET. First, we explore *context-aware particle picking* as a method to retain biological context when selecting particles for subtomogram averaging and other downstream analyses. Secondly, we demonstrate *area-selective template matching*, where we use segmentation-based masks to avoid redundant computations in template matching and enable >500-fold faster processing in specific cases. To illustrate the utility of the approach, all segmentation results have also been made available online via cryopom.streamlit.app.

## Introduction

Cryo-electron tomography (cryoET) combined with cryo-focused ion beam (cryoFIB) sample preparation enables the visualization of the structural biology of cells and tissues^1–4^. Recent advances in instrumentation and methodology, such as the automation of lamella preparation^5–8^, fast tilt-series acquisition^9–11^, and unsupervised tomogram reconstruction methods^12–14^, have enabled the acquisition of data at a much faster rate than was previously possible. As a result, the field is now poised to achieve large-scale quantitative studies cryoET^15,16^: generating sufficiently large datasets to harness not only on the high resolution provided by cryo-electron microscopy, but also the high context view of the cell that is unique to cellular cryoET^17–19^.

As datasets scale, scalable data analysis strategies are also required^20–23^. Whereas it was previously often feasible to select particles or subsets of data manually, the management of larger datasets requires alternative methods of data curation that can be applied at scale. This requirement extends to downstream cryoET processing tasks such as segmentation, particle picking, and subtomogram averaging, where current unsupervised methods struggle with large data volumes or contextual preservation^24–29^.

Aside from introducing technical challenges, the ability to perform large-scale studies with cryoET also introduces new possibilities for including more information about biological context during high-resolution structure determination. Thus far, protein structure determination by subtomogram averaging has largely been a 3D-variant of single-particle cryoEM, where the structures of proteins are determined at high resolution but in the absence of any biological surrounding: after locating particles in tomograms, potentially relevant context is largely discarded as particles are represented only as coordinates, which are then used to extract particles and provide the input for the subtomogram averaging pipeline^30^. Since the major benefit of performing cryoET over single-particle cryoEM lies in its ability to visualize structures in their native biological context, making use of this context in downstream processes will enable more nuanced interpretations of the resulting structures.

Here, we explore a comprehensive segmentation strategy as a tool for large scale studies with cryoET. To demonstrate the strategy we use the dataset by Kelley and Khavnegar et al.^15,16^, which comprises 1829 tomograms acquired on cryoFIB-milled lamellae of the alga *Chlamydomonas reinhardtii* (CryoET Data Portal^31^ IDs DS-10302, DS-10301). Since the internal structure of this photosynthetic unicellular eukaryote is highly complex and comprises a large variety of subcellular features^32,33^ and the data is of exceptional quality, this dataset provides an ideal environment for testing analysis methods in cryoET^16^.

By segmenting over twenty distinct ontology-based subcellular features, including both macromolecular complexes and organelles, we show how a large dataset can be ‘summarized’ and represented as a searchable dataset, enabling data curation as well as discovery within the data. Building upon this strategy, we propose two applications of ontology-based segmentations in improving the throughput or context-awareness of common cryoET processing tasks. We think that both applications could be of interest specifically in the context of increasingly unsupervised analyses on ever-growing datasets.

First, we demonstrate *context-aware particle picking* as a way of integrating information about the biological context of particles in the particle picking and subtomogram averaging workflow. Using ribosomes, ATP synthase, and RuBisCo as exemplary targets, we show how the use of segmentation-derived *context-vectors* – i.e. numerical representations of the subcellular environment within which a particle was detected – enables curation and classification of particles based on biologically relevant parameters.

Secondly, we propose *area-selective template matching* as a way to aid scaling up applications of template matching (TM). By combining different segmentations and based on the prior information that nuclear pore complexes only occur within the NE, we show that the usage of segmentation-based masks that limit the application of TM to areas that are likely to contain a structure of interest is a promising approach for increasing the throughput of TM-based approaches to particle detection.

To accompany the results outlined in this article, all segmentation results are also available online via the CryoET Data Portal^31^ under deposition ID 10314, or can be explored in the automatically generated segmentation report hosted at cryopom.streamlit.app. The open-source software that was developed to enable these experiments and visualizations is available online via github.com/bionanopatterning/Pom or on the Python package index as *Pom-cryoET*.

## Results

### Segmentation strategy

Whereas segmenting organelles such as nuclei or mitochondria is a common application in the field of volume electron microscopy^34,35^, the scale of cryoET is typically such that macromolecules themselves, rather than the larger structures they constitute, are the foremost features of interest for segmentation^20,36–39^. Membranes, ribosomes, and ice particles, for example, form very typical patterns that convolutional neural networks are very efficient at recognizing and annotating^40–42^.

Organelles, in comparison, are much more varied in appearance and shape, and manifest in images as much larger-scale patterns. As a result, segmenting organelles in cryoET datasets can be relatively challenging^25^. Yet, the accurate segmentation organelles types could make a meaningful contribution to the biological interpretation of data: it presents information about the local cellular context, as well as narrows down the subset of macromolecular structures that may be found in an area.

Although the molecular composition of cellular tomograms is very complex, highlighting a select few macromolecular structures can already provide enough information to enable the identificiation of a large number of different subcellular environments. For example, even if we could only observe membranes, we might already distinguish mitochondria, the Golgi apparatus, and the nuclear envelope, each by characteristically shaped membrane. Including ribosomes in the observation, one could also differentiate between the cytoplasm and nucleoplasm, and perhaps between the rough and smooth endoplasmic reticulum.

To bridge the different scales of macromolecules and organelles, we tested a two-step segmentation approach that was based on this notion, involving the sequential application of two convolutional neural network (CNN)-based segmentation methods: first, segmenting ***density to macromolecules,*** and second, segmenting ***macromolecules to organelles***. By using macromolecule segmentations as an intermediate representation, we specifically aimed to strike a balance between accuracy and practicality: the two separate segmentation procedures are relatively straightforward for a user to prepare, and the network architectures that we use are small enough to enable high-throughput parallel processing.

The first step in this workflow was to segment density into macromolecules. Among various clearly recognizable macromolecules visible in the *C. reinhardtii* dataset, we decided to limit our selection to **membranes**, **ribosomes**, (mitochondrial) **ATP synthase**, and **RuBisCo** (**Fig. 1AB**). The motivation to limit the selection to just these four structures was twofold: first, they are among the most prevalent and easiest to annotate structures visible in the tomograms. Secondly, we expected that this set would usefully represent the distinct molecular compositions of the various organelles we would aim to segment (see below). In order to perform these automated segmentations efficiently we made a number of improvements to the cryoET segmentation tool Ais^43^ (**Fig. 1 supplement 1**), which enabled us to segment all features at a rate of approximately 1 second per macromolecule type per tomogram, or about 2 hours of processing in total for the 1829 tomograms (see Methods). The improvements included the implementation of significantly faster parallel processing on multiple GPUs and additional features for organising many-feature annotations.

**Figure 1.**
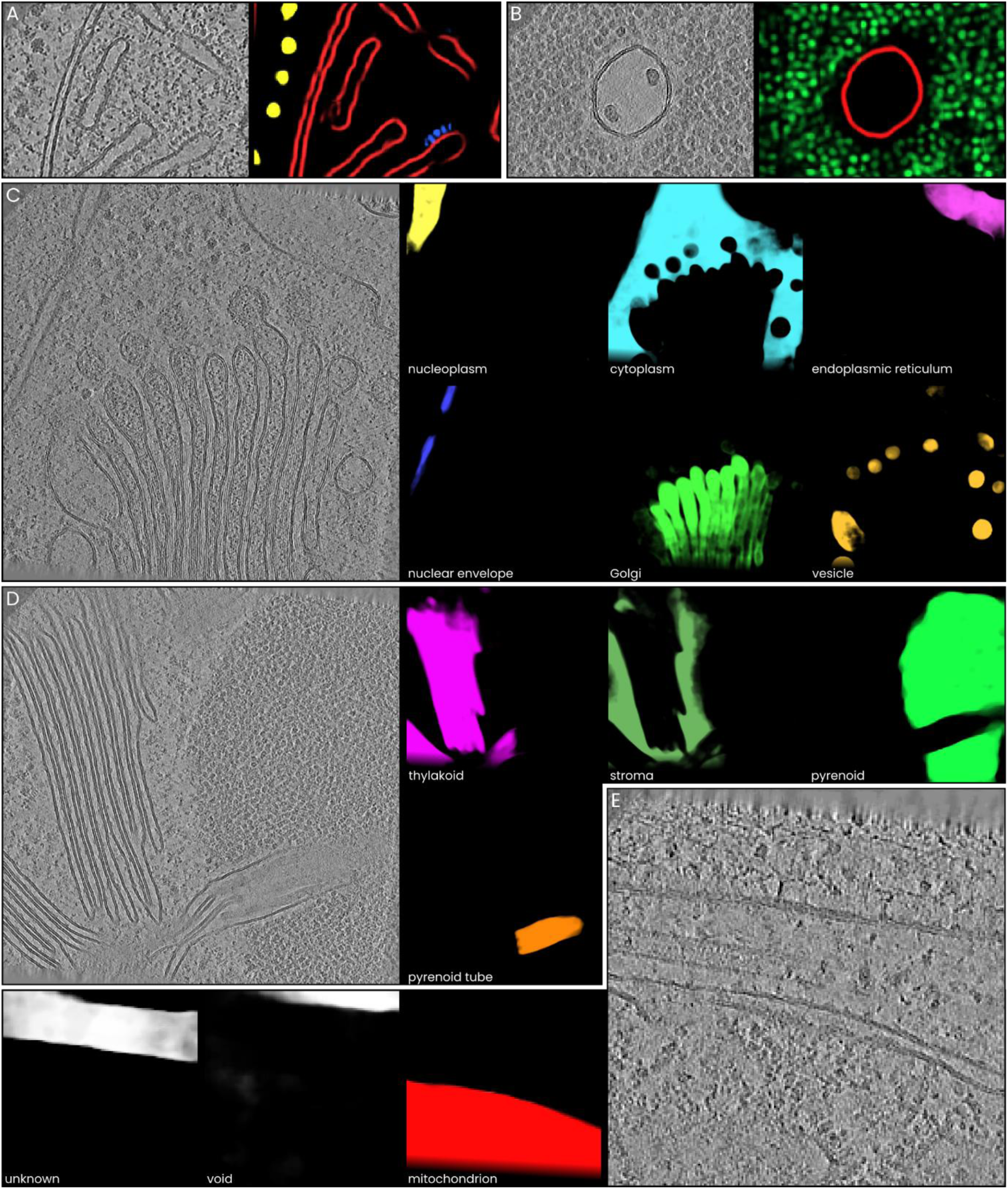
An overview of the segmented macromolecules and organelles. **A)** A small region of a tomogram containing a mitochondrion, with ribosomes (yellow), membranes (red), and ATP synthase (blue) visible. **B)** A small region of a tomogram containing pyrenoid and pyrenoid tubules, with membranes (red) and a large number of RuBisCo (green) molecules visible. **C)** A tomogram acquired close to the nucleus, showing nucleoplasm (yellow), nuclear envelope (blue), cytoplasm (cyan), Golgi (green), endoplamis reticulum (magenta), and various vesicles (amber). **D)** A tomogram that features various photosynthesis-related organelles: the thylakoid (magenta) and stroma (olive) of the chloroplast, as well as pyrenoid (green) and a pyrenoid tubule (orange), which are also called thylakoid tubules, since these structures are continuous with the thylakoid membranes of the chloroplast. **E)** A small region of a tomogram featuring a section of mitochondrion (red) as well as the two special classes: void (corresponding, i.a., to the reconstruction artefact in the top-right corner) and unknown (a ‘leftover’ class, covering all image regions where the neural network does not identify any of the other classes)

The second step was to segment different organelles and subcellular regions, using the density and macromolecular information as the input data. Our selection of features (see Methods for corresponding Gene Ontology IDs^44^) comprised the **cytoplasm**, **nucleoplasm**, **nuclear envelope**, **endoplasmic reticulum**, **mitochondrion**, **Golgi**, **vesicle**, **thylakoid**, **stroma**, **pyrenoid**, **pyrenoid tube**, and **void** and **unknown**, the latter two being special cases that we discuss below. Achieving automated segmentation of these features required a number of steps, which are described in more detail in the **Supplementary Information**. In brief, we first trained multiple separate CNNs that each segmented one class using manually-annotated images as the training data. We then applied each of these preliminary single-feature CNNs to every separate training dataset, and combined all the resulting segmentations (without further curation) to prepare a single training dataset for use in training a final network that could segment all classes simultaneously. The resulting network could segment the 13 output features within all 1829 tomograms in less than one hour (see Methods).

### Ontology-based summarization and curation of datasets

A relatively straightforward application of this segmentation approach was then to use the resulting volumes to ‘summarize’ the dataset, so that instead of represented as thousands of separate volumes, it could be represented in a condensed human- and machine-readable format. One possible way of achieving this is simply by measuring the make-up of each individual tomogram in terms of its fractional composition: percentage values that indicate what fraction of the tomogram volume corresponds to cytoplasm, nucleoplasm, or any of the other segmented classes. For example the composition of a single tomogram (**Fig. 2A, B**) could be summarized as comprising 26% Golgi, 22% cytoplasm, 8% nucleoplasm, 2% nuclear envelope, 2% vesicle, and the remainder mostly void.

**Figure 2.**
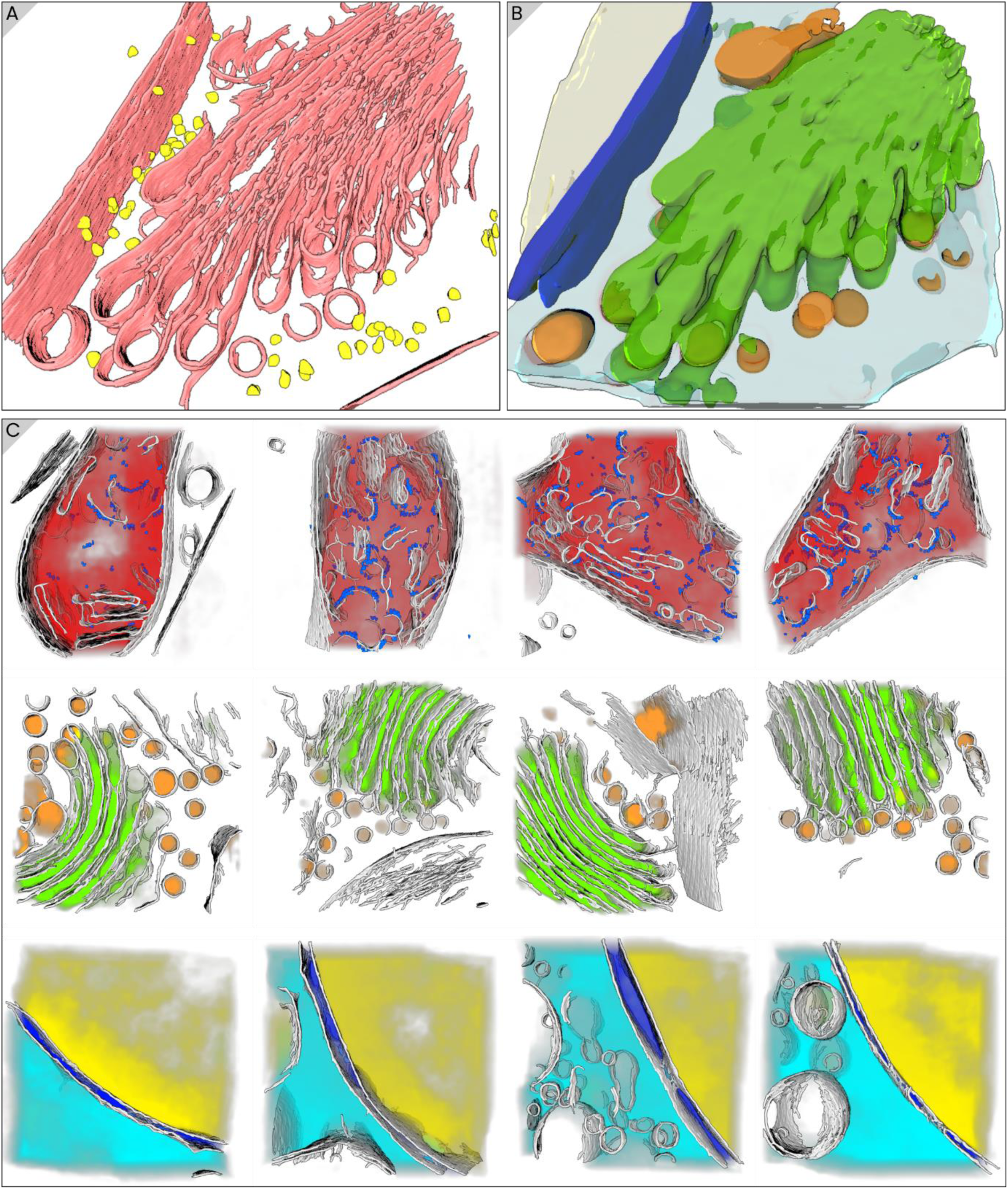
Exemplary segmentations of curated sub-datasets. **A)** Macromolecule (membranes in red, ribosomes in yellow) segmentation, and **B)** organelle segmentation of a tomogram featuring Golgi (green), vesicles (orange), cytoplasm (cyan), nucleoplasm (beige), and nuclear envelope (blue). **C)** Examples of sub-datasets, selected by filtering the data for certain criteria. Top: mitochondrion-containing tomograms, showing mitochondrion (red), membrane (gray), and ATP synthase segmentations (blue). Middle: tomograms that contained significant volumes of the Golgi apparatus (green) and associated vesicles (orange). Bottom: tomograms containing 10% or more cytoplasm (cyan) as well as nucleoplasm (yellow). The nuclear envelope (blue) is also rendered. In this panel, macromolecule segmentations are rendered as isosurfaces and organelle segmentations as emissive volumes with intensity proportional to the segmentation value.

By condensing and tabulating the composition of all tomograms in the dataset in this manner, segmentation results can be used to summarize data in a format that enables straightforward curation. For instance, it enables the selection of data subsets (**Fig. 2C**), such as ‘tomograms with the largest mitochondrion content’ (**Fig. 2 supplement 1**), ‘tomograms containing at least 10% Golgi’ (**Fig. 2 supplement 2**), or ‘volumes containing at least 10% cytoplasm and nucleoplasm’ (**Fig. 2 supplement 3**). By combining multiple filters, it is also possible to search for specific, possibly rare tomogram compositions. For example, although many tomograms contain either >5% thylakoid (n = 651) or >5% pyrenoid (n = 74), only a small subset (n = 7) of tomograms feature both classes; yet, these tomograms could be of particular interest for studying interactions between chloroplasts and pyrenoids (**Fig. 2 supplement 4**). An interactive summary of this dataset is available online at cryopom.streamlit.app.

### Achieving a comprehensive set of output classes

In the introduction we mentioned that besides a number of biological features, the network also output two special classes: **void** and **unknown.** The inclusion of these two classes serves to achieve two goals.

First, the annotations for the **void** class were prepared in such a manner that the void output roughly corresponds to areas of ‘low data quality’ (**Fig. 3 supplement 1**). Such areas included those featuring padding artefacts in the tomograms, poorly reconstructed volumes, regions of the reconstructed volume that lie above and below the edges of the lamella, and regions where large ice particles contaminated the view. In total ∼43% of the data was classified as void (mostly because the reconstruction volume was larger than the actual sample was thick).

**Figure 3.**
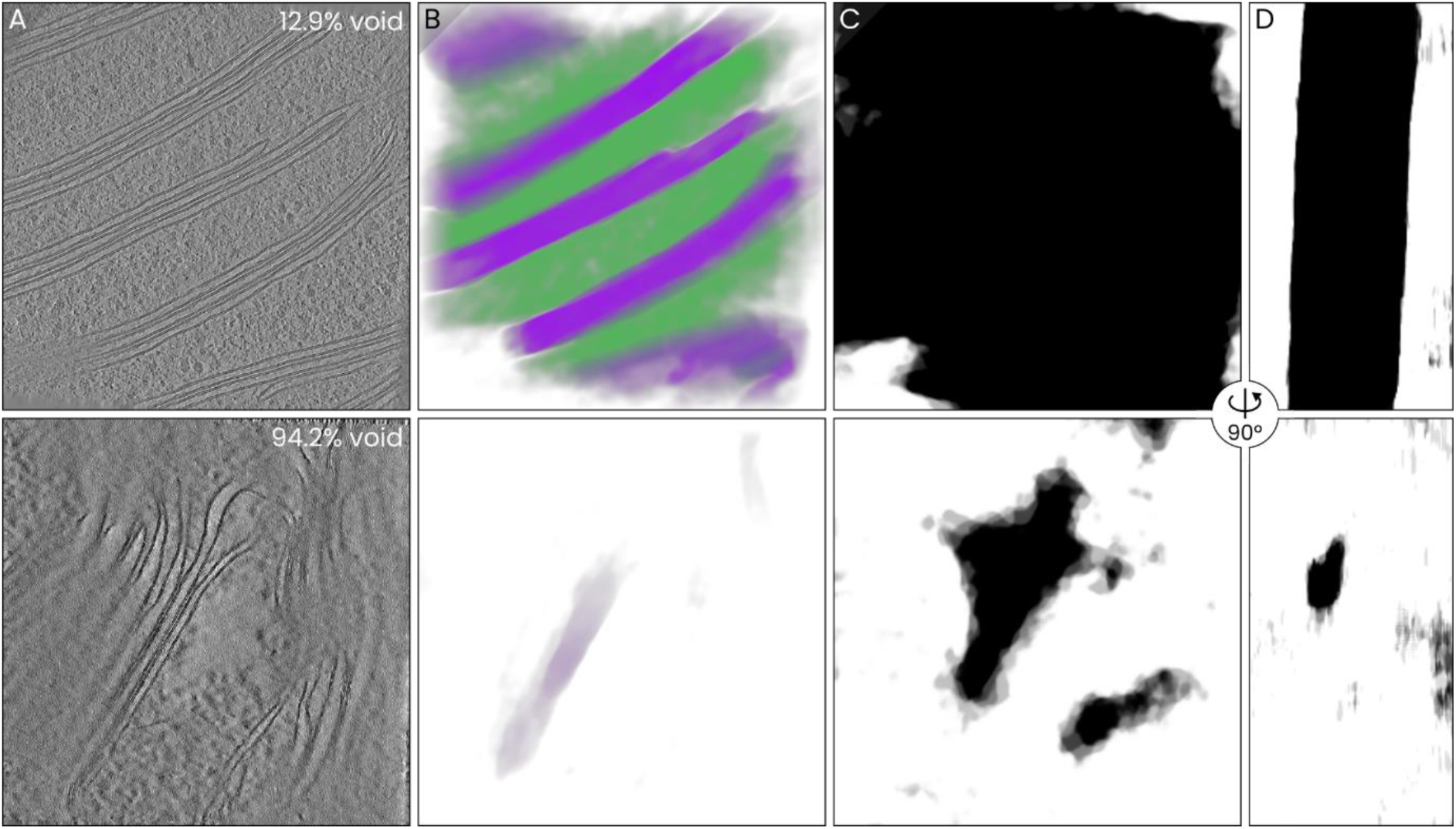
The void output class is a metric for tomogram quality and covers a large fraction of the total data volume. **A)** The central slices of two tomograms, both manually identified as containing mainly thylakoid and stroma, with low and high void volume fractions. A value of 12.9% (top) is in the top-50 lowest values, while a value of 94.2% (bottom) is among the highest. The difference in data quality is apparent. **B)** Corresponding thylakoid (purple) and stroma (green) segmentations. **C)** Corresponding void segmentation output for the slices shown in A. **D)** Corresponding void segmentations for central YZ slices. When the tomogram quality itself is high (top), the outline of the lamella’s volume can be recognized in the void segmentation.

As a result, the void output can be used as a rough metric of tomogram quality (**Fig. 3**). Tomograms for which the mean void output value is low are typically well-reconstructed tomograms with easily identifiable and segmentable macromolecular features (**Fig. 3A, B**). Tomograms with a high void value are typically of relatively low quality; often due to a high lamella thickness or poor tilt series alignment. Within individual volumes, the void class also serves to differentiate between low and high quality image regions (**Fig. 3C**) and, when the tomogram quality is high, it can be indicative of the boundaries of the lamella within the reconstructed volume (**Fig. 3D**).

Secondly, the “**unknown**” class is used as a ‘leftover’ class to annotate those features that are of sufficient quality (as measured by the void output), but that do not correspond to any of the selected biological output classes. Since the neural network used for the organelle segmentation uses softmax activation in the final layer (ensuring that the sum of all different segmentation values for every pixel equals one, and thus that the output can be interpreted as a probability distribution), including an extra class for unspecified features is necessary to ensure the normalization does not arbitrarily scale the output of other classes.

By providing the neural network with training data for the unknown class that corresponds to regions where all intermediate single-feature models output a low value, the resulting ‘unknown’ segmentation output corresponds to image features that were both of high quality (as ensured by the void class) and that were not included in the selection of biological classes. As such, the unknown class can be used to identify biological features that should be included in the ontology, in order to comprehensively represent the features present in the data.

In the present case, around 5% of the data was classified as ‘unknown’ (versus 52% that was covered by the 11 biological classes and 43% by void). While some of this output could be considered artefactual (**Fig. 4 supplement 1**), it also included a number of specific biological features such as: a) **lipid droplets,** b) **nuclear pore complexes,** c) the **cell wall,** d) **electron dense layers, and** e) **intermediate-filament-rich areas** (**Fig. 4**).

**Figure 4.**
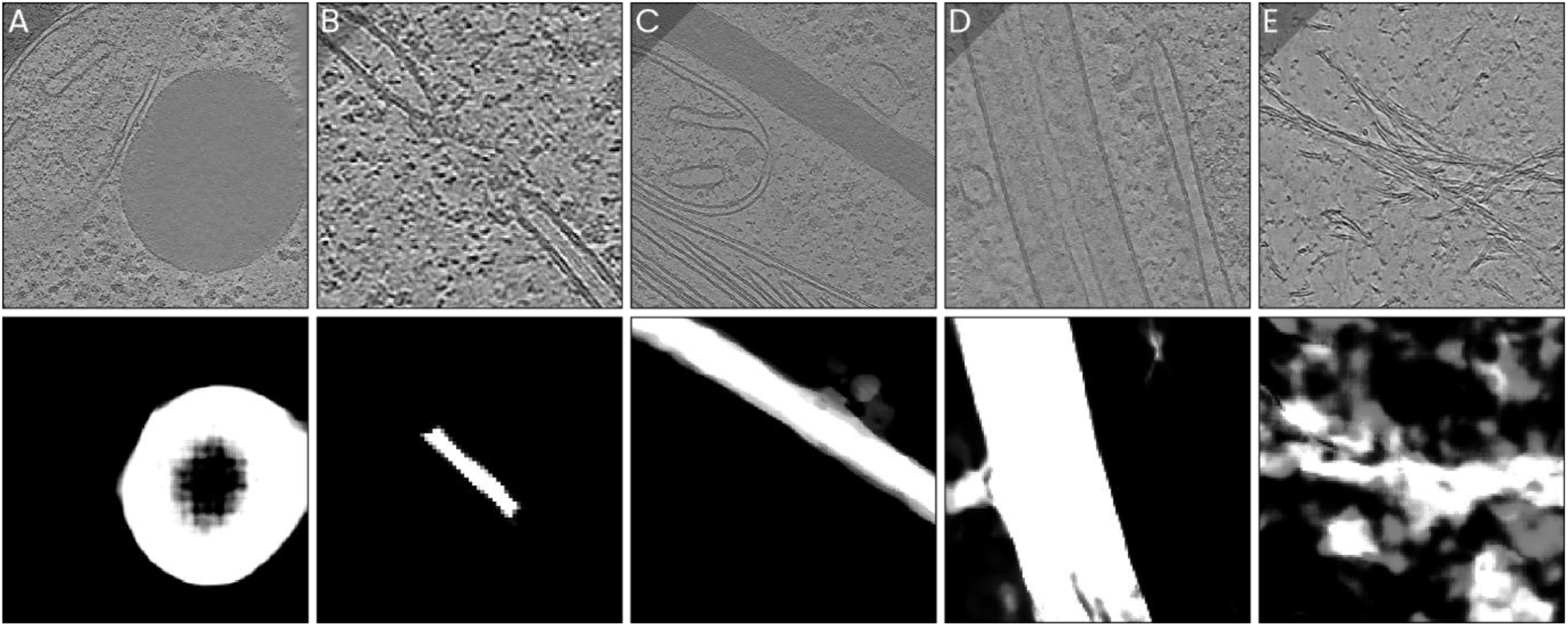
Five distinct subcellular features that are annotated as ‘unknown’. The void output class is a metric for tomogram quality and covers a large fraction of the total data volume. Density (top) and corresponding unknown class outputs (bottom) are shown adjacently. **A)** A lipid droplet, **B)** a nuclear pore complex, **C)** a thick electron-dense layer, **D)** the cell wall (of adjacent cells, in this example), **E)** intermediate-filament-rich areas.

On the basis of these results, we decided to additionally include lipid droplets, nuclear pore complexes (NPCs), and the dense layer in our selection of segmented features. After adding annotations for these classes, training a new iteration of the network, and processing all volumes, we could then use the results to specifically find additional instances of NPCs, LDs and DLs in the data (**Fig. 4 supplements 2-4**). In a third and final iteration, we also included the **chloroplast outer membrane**, **cilium, cell wall, starch granules**, and **intermediate-filament-rich areas (IFRA)** in the segmentation (**Fig. 4 supplements 5-9**), bringing the number of segmented features up to 23 (excluding void and unknown). At this point, the remaining unknown-annotations were mostly of the aforementioned artefactual nature.

### Context-aware particle picking

Particle picking in cryoET involves the identification of specific particles within volumes and listing their coordinates. These coordinates can then be used as the input for subsequent processing steps, such as *subtomogram averaging* (STA). By extracting a particle from its environment, information about the particle’s relation to other structures within the cell can be lost.

Ribosomes occur in many different types that are found in distinct cellular environments. The 80S ribosome, for instance, can be found bound to the endoplasmic reticulum (ER) or to the nuclear envelope (NE), or in the cytoplasm. Subunits of the 80S ribosome can also be found in the nucleus, where they are synthesized^45^, while additional distinct types of ribosomes also occur in mitochondria and chloroplasts. Despite differences in structure, these different types of ribosomes can be difficult to differentiate in cryoET data^46,47^.

Comprehensive segmentation offers a way of including the local cellular context of a particle in the picking step. Instead of considering only a particle’s coordinate, one could approximately ‘measure’ the particle’s cellular context by sampling all segmented features in a small region of interest surrounding the particle. In the case of ribosomes, for instance, measuring the average values of the different segmentations within a ∼20 nm radius from a ribosome coordinate assesses whether the ribosome is in close proximity to the ER, NE, or to a membrane, whether it is surrounded mainly by cytoplasm, nucleoplasm, or stroma, or whether it is found within a mitochondrion.

To test this *context-aware particle picking* (CAPP) approach, we used the previously generated ribosome segmentations as a basis to automatically pick ribosomes in all 1829 tomograms (**Fig. 5A**, see Methods). Next, we sampled all organelle segmentations as well as the membrane segmentation in an approximately 25 × 25 × 25 nm box centered at each ribosome coordinate (**Fig. 5B**), and stored the average segmentation value in a list that we called a *context vector*. The resulting particle dataset comprised a little under 300,000 ribosome coordinates, as well as an equal number of corresponding 15-valued context vectors.

**Figure 5.**
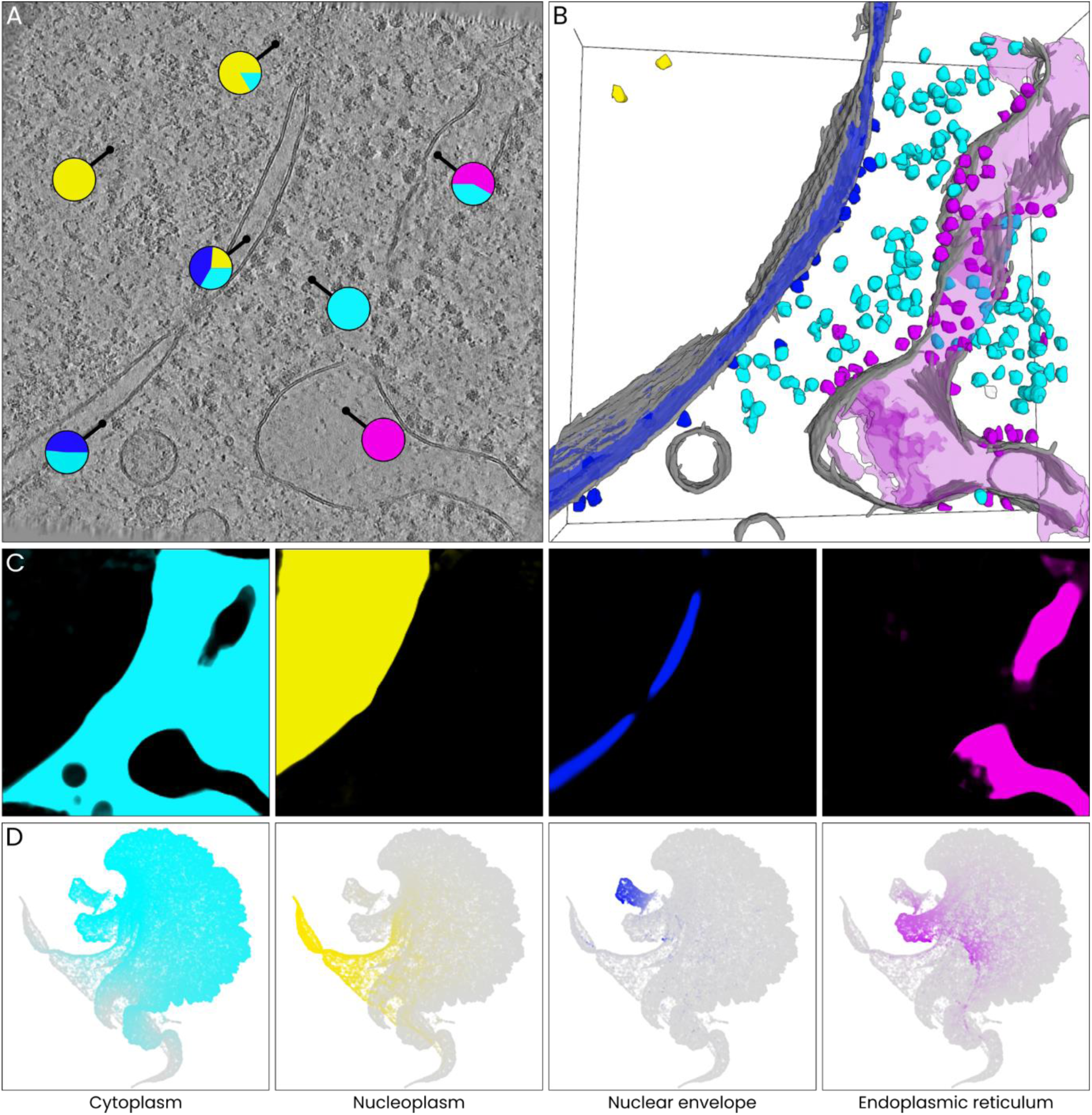
Context-aware particle picking applied to ribosomes. **A)** A schematic representation of the meaning of ‘context vectors’. By sampling all organelle segmentations in small areas of interest within a volume, distinct cellular environments can be identified. Here, local contexts are illustrated as pie charts. Aside from definite nuclear (yellow), cytoplasmic (cyan), and ER (magenta) areas, mixed contexts are measured at interfaces between different subcellular environments; for example, the context vector for a ribosome bound to the NE would have a high value for both cytoplasm and NE, and possibly also for nucleoplasm (depending on the size of the volume within which the context is sampled). **B)** A 3D visualization of the membrane (gray), NE (blue), ER (magenta), and ribosome segmentations for the same tomogram as in A. Each ribosome particle was assigned a colour based on the corresponding vector (magenta if >10% ER, blue if >10% NE, yellow if >30% nucleoplasm, cyan if >50% cytoplasm, else white) **C)** From left to right: the cytoplasm, nucleus, NE, and ER segmentations for the slice shown in A. **D)** UMAP of all context vectors, with points coloured by the different relevant elements of the context vector (labelled below).

To assess whether these context vectors reflected the cellular context within which a particle was detected in a useful way we generated a 2D UMAP representation of the context vectors in order to visualize how the particles cluster in the context space. We then highlighted those cellular features that are relevant to subclasses of 80S ribosomes: nucleoplasm, cytoplasm, NE, and ER (**Fig. 5C**).

As expected, the resulting map (**Fig. 5D**) demonstrated a number aspects of the distribution of ribosomes. First, it shows that a majority of the detected particles corresponded to free cytoplasmic ribosomes. Secondly, distinct groups corresponding to nuclear, NE-, and ER-associated ribosomes were also visible. Additional subgroups, such as mitochondrion and stroma-associated ribosomes could also be identified (**Fig. 5 supplement 1**).

Finally, the ‘void’ context value was found to be high for a relatively large number of ribosomes. By inspecting cases where the void score was low or high, it seemed likely that the void value could be a useful metric to discard spuriously picked particles from the selection (**Fig. 5 supplement 2**). Results of similar tests of CAPP, focusing on ATP synthase (**Fig. 5 supplement 3**) and RuBisCo (**Fig. 5 supplement 4**), suggested that the usage of context-vectors in curating particle selections might be a generally useful strategy.

### Area-selective template matching

Like segmentation-based approaches, template matching (TM) is a strategy to automate the detection of specific structures within tomograms. Despite recent advances in software^27,48,49^, TM remains a very computationally demanding task. As a result, and especially when datasets are large and the structures of interest are small, TM is currently not often a practical approach. Yet, it is a promising method for use in unsupervised data mining strategies, where TM might be a useful and relatively unbiased approach to detect structures that are rare, small, or otherwise difficult to detect manually.

Since specific structures of interest invariably occur only within specific subcellular environments, ontology-based segmentations can offer a way of incorporating prior biological information into an analysis in order to increase the throughput of TM. For instance, when the targets of interest are microtubules, limiting the area of interest to the cytoplasm could be a practical way of reducing the computational load: only ∼18% of the data volume comprises cytoplasm. In some cases, the volume of interest may be even lower; for instance, if NE-bound ribosomes are the target, limiting the search to the ∼0.4% of the data volume that corresponded to NE might help reduce the computational cost by over two orders of magnitude.

Nuclear pore complexes (NPCs) (**Fig. 6**) are a good example to demonstrate *area-selective template matching* (ASTM) as a particle detection strategy that combines the efficiency of CNN-based segmentation and the more specific and less supervised method of TM. Although one could also pick NPCs on the basis of segmentation results, detecting particles via TM instead has two advantages: first, TM should be better able to avoid false positives, as it is less prone to bias than machine learning-based methods. Secondly, TM yields not just particle coordinates, but also orientations, which are an important aspect of NPCs.

**Figure 6.**
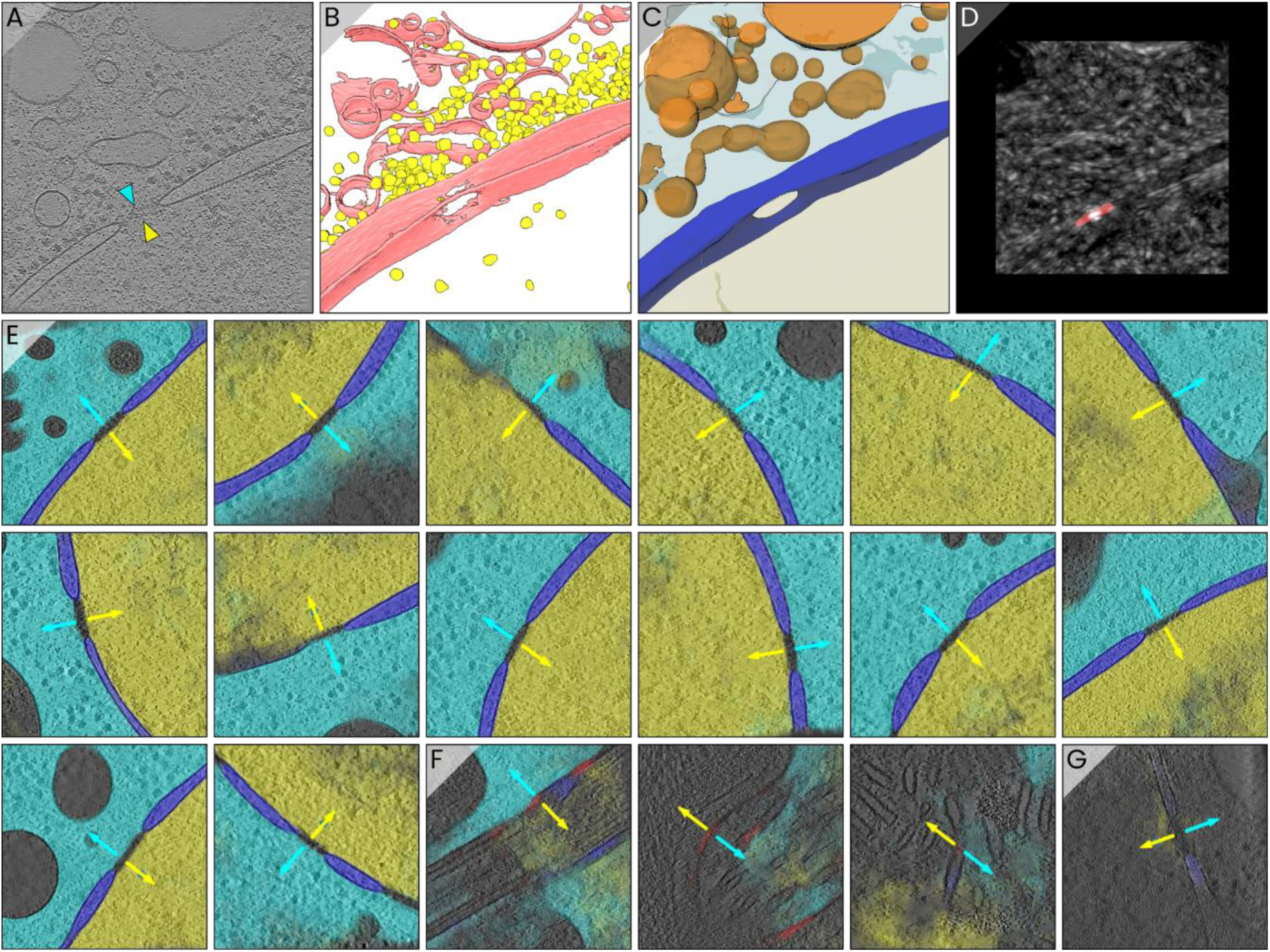
Area-selective template matching detection of NPCs. **A)** The nuclear pore complex used as the template for ASTM. The detected cytoplasmic (cyan) and nuclear (yellow) sides are indicated by markers. **B)** Macromolecule segmentation (membranes in red, ribosomes in yellow) of the tomogram in A, **C)** Organelle segmentation, displaying nucleoplasm (beige), nuclear envelope (blue), cytoplasm (cyan), and vesicles (orange). **D)** A map of the TM scores for the full slice as displayed in A. The NPC-segmentation derived mask that would be used for ASTM is overlayed in red, in the location where a high TM score was indeed computed. **E)** A collage of automatically picked and oriented NPCs. Cytoplasm, nucleoplasm, and nuclear envelope segmentations for the same slice are overlayed in cyan, yellow, and blue, respectively. **F)** Examples of false positive identifications. In each of the three panels the NPC-segmentation is overlayed in red. Where both the NPC-segmentation output false labels and the local density resembles the NPC template, ASTM picking yields false positives (as would TM). **G)** An exemplary case where an NPC was correctly identified despite the image quality being relatively low and the cytoplasm, nucleoplasm, and nuclear envelope segmentation outputs failing.

Using a custom GPU-accelerated 2D real-space TM implementation capable of area-selective processing (see Methods), we applied ASTM to locate NPCs in all 1829 tomograms in the dataset. We manually sampled a single NPC from one of the tomograms for use as the template (**Fig. 6A-C**), and thresholded the NPC-segmentations at half the maximum output value to generate the required masks (**Fig. 6D**). Using a single GPU (NVIDIA P2000), the template matching step itself took on average around 0.6 seconds per tomogram with ASTM versus 395 seconds with conventional TM. The use of segmentation-bases masks thus sped up the computation more than 500-fold in this case.

When the processing was completed, we had identified 385 NPCs across the dataset. By sampling the cytoplasm and nucleoplasm segmentation results on both sides of each pore, we then also refined the pore’s orientation so that each NPC could be represented as an oriented point, with defined cytoplasmic and nuclear sides (**Fig. 6E**). Upon manual inspection, 17 particles (4.4%) were identified as false positives. These occurred in locations where both the neural network output was erroneous (labelling, e.g., cristae in a mitochondrion as NPC) and the local density resembled that of NPCs (**Fig. 6F**). The number of false negatives is harder to assess, but since the NPC-segmentation output accurately annotated NPCs even in cases where segmentation results for other classes failed (**Fig. 6G**), we deem it likely that possible false negatives would mainly occur in tomograms of relatively low quality. A second test, targeting mitochondrial ‘lightbulb’ complexes (see Khavnekar et al. (2023)^15^ and Waltz et al. (2024)^50^), yielded similar results: the use of area-selective masks to perform TM only within a ∼20 thick shell around mitochondria sped up the computation by over >50 fold, and yielded a small set of additional lightbulb particles based on a single observation of its kind (**Fig. 6 supplement 1**). Although the accuracy of these lightbulb detections could be much improved and the throughput could also be substantially higher if parallel computation on multiple computers or GPUs was employed, our results suggest that ASTM can be a useful strategy for scaling the application of TM towards larger datasets.

## Discussion

In this article we proposed and demonstrated an iterative workflow to achieve comprehensive – i.e., assigning identities to all voxels in a dataset – segmentation of large cryoET datasets. Using a set of 1829 tomograms by Kelley and Khavnegar et al., we segmented **membranes, ribosomes, ATP synthase** and **RuBisCo** as macromolecular structures, and **cytoplasm, mitochondria, nucleoplasm, nuclear envelope, nuclear pore complexes, pyrenoid, pyrenoid tubes, thylakoid, stroma, Golgi, vesicles, endoplasmic reticulum, lipid droplets,** and an unidentified **dense layer** as organelles, plus **void** as a data quality metric and **unknown** as a means of highlighting the remaining unidentified cellular features. Two further macromolecule types, **proteasomes** and **T-complex protein ring complexes (TRiC)** and five additional organelle-like features, **starch granules**, **cilia**, **intermediate-filament-rich areas (IFRA), cell wall,** and **chloroplast outer membrane**, were added to the segmentation in subsequent iterations of the workflow. The resulting 49383 segmentation volumes are available on the CZII CryoET Data Portal under deposition number 10314.

On the basis of this workflow we then proposed two data mining strategies that may be useful when working with large datasets. The first of these, **context-aware particle picking** (CAPP), was suggested as a tool to curate particle selections, as well as a method by which to incorporate details about particles’ cellular environment in downstream cryoET analyses such as subtomogram averaging. Although we did not show these applications yet (lacking CTF parameters and the original unbinned tilt series, we decided against testing these for the time being), it would be interesting to see the extent to which the proposed context vectors can be used to inform heterogenous reconstructions, or vice versa, in large scale cellular cryoET studies. Secondly, we proposed **area-selective template matching** (ASTM) as a strategy to combine the efficiency of automated segmentation with the improved specificity and reduced bias of TM.

Exploring the potential of CAPP and ASTM would require much further testing. For example, it could be interesting to combine CAPP with heterogenous reconstruction, in order to investigate whether context-vectors correlate with particle embeddings in the latent configuration space, or use it for classification in homogenous subtomogram averaging. In preliminary experiments, we found that subtomogram averaging (STA) of subsets of ribosomes did clearly result in membrane-bound structures for the NE- and ER-associated particles, while cytoplasmic ribosomes appeared not to be. However, with our data limited to a resolution of 15.68 A /pixel (originally 7.48 A /pixel on the data portal) and without access to the original tilt series and CTF-parameters, the achievable resolution in STA was limited, and we decided to limit our current scope and did not pursue further, more thorough STA analyses.

Likewise, although we demonstrated the feasibility of ASTM for NPC detection, both the implementation of ASTM and its general applicability for targeting other complexes require further refinement and testing. Implementing ASTM efficiently is certainly non-trivial, and the utility of segmentation masks importantly also depends on whether real-space or Fourier-space TM is employed. In the latter case, masks could in principle be used to crop smaller regions of interest from a tomogram, but pixel-wise masking with the goal of reducing computational load is not a straightforward option due to the nonlocal nature of Fourier-space TM. Nevertheless, combining segmentation-based selection of small data-subsets with conventional TM could be a useful strategy to increase processing throughput.

A number of notable limitations to our segmentation approach could be significantly improved upon. First, although the VGGNet and UNet neural network architectures that we used are efficient both in training and application, more advanced architectures would likely enable better performance^25^. Specifically for the organelle segmentations, the accuracy was noticeably hampered by the limited size of the UNet’s receptive field. While the global context of a tomogram provides much information about local feature identities, the UNet was not able to make use of it. Secondly, manual annotation of the training data remains a laborious task, especially when the number of features to segment grows larger. Incorporating additional strategies for training data augmentation or unsupervised learning to lower the time cost of labelling would be a useful improvement. Third, although the concept of meta-segmentation (segmenting organelles using macromolecule segmentations as the input) appeared to improve the segmentation accuracy at least marginally, more sophisticated networks or networks provided with more comprehensive training inputs could likely achieve direct segmentation from densities to organelles with the same or better accuracy and processivity; because if a combination of networks can do it, then so can a single network.

Another observation that we think is important relates to the availability of computer infrastructure. The simple reason why we could segment all 1829 volumes in less than one hour (for the organelle segmentations) was that we had access to four Linux servers that were equipped with 8 GPUs each – had we been limited to using a single GPU, the process would have taken almost 1.5 days. In our experience, an important practical requirement for generating accurate CNNs is the ability to iterate: to test a network, expand or improve the training data as required, and then test again. This is especially the case when a large fraction of the data is never seen prior to applying the networks. In such a case, processing all volumes and exploring the output in an interactive way, as enabled by the cryopom.streamlit.app interface, can be a very useful way to identify biases and flaws in the network output. Such iterative and interactive refinement requires sufficient computational resources. As datasets continue to grow, the importance of investing in accessible computing infrastructure is thus likely to grow along.

Finally, this work would not have been possible without the aid of initiatives like EMDB, EMPIAR, and the CZII CryoET Data Portal, and the researchers who contribute data to these platforms. The high cost of cryoET hardware and infrastructure are currently significant barriers for many research groups, and the preparation of quality samples one of the major bottlenecks in cryoET studies. The free availability of data is therefore essential not just for the development of machine learning methods or particle picking and reconstruction algorithms, but also for enabling biological discovery. This is especially the case as the field moves forward into quantitative cryoET. As such, to ensure that researchers worldwide can benefit from the rapid advances in cryoET, it is critical that we engage in collaborative and open sharing of data.

## Methods

### Computing setup

After preparing annotations on a desktop Windows PC, all training and processing tasks were conducted on four remote Linux (Debian) servers, each equipped with eight CUDA-enabled NVIDIA GPUs each (RTX 3080 and RTX 2080 Ti). Training required typically around 15 minutes for the Ais models, while training for the intermediary single-feature organelle networks ranged from 10 to 45 minutes, depending on the size of the training dataset. For the organelle-segmenting networks, we eventually settled on 100 epochs of training for the single feature network and 300 for the shared network. Training for the latter took around 3 hours.

We used slightly different data-parallel approaches for training and for inference. During training we used TensorFlow’s mirrored distribution strategy, where the model is replicated on all GPUs and batches of data are processed in parallel. For inference, we launched eight separate asynchronous threads on each server, each thread using a single GPU to process a subset of all tomograms. One thread required on average around 60 seconds to process a single volume using the shared model, so that processing the full dataset (1829 tomograms) required just under one hour when using all 32 GPUs. It is important to note that the high throughput of our approach is thus primarily due to the availability of a large number of GPUs, which allows for efficient parallel processing and significantly reduces the overall processing time.

### Training data and feature definitions

The number of training images used for each of the various single-feature networks (macromolecules and intermediary organelle-segmenting networks) summarized in the table below. Negative examples are images that do not show the feature of interest and that are not annotated. The total manually annotated image area was equivalent to ∼1.5 full tomograms, or 0.083% of the full dataset. The rightmost columns, ‘**GO ID**’, lists the corresponding Gene Ontology^44^ ID for each feature.

**Table 1.**
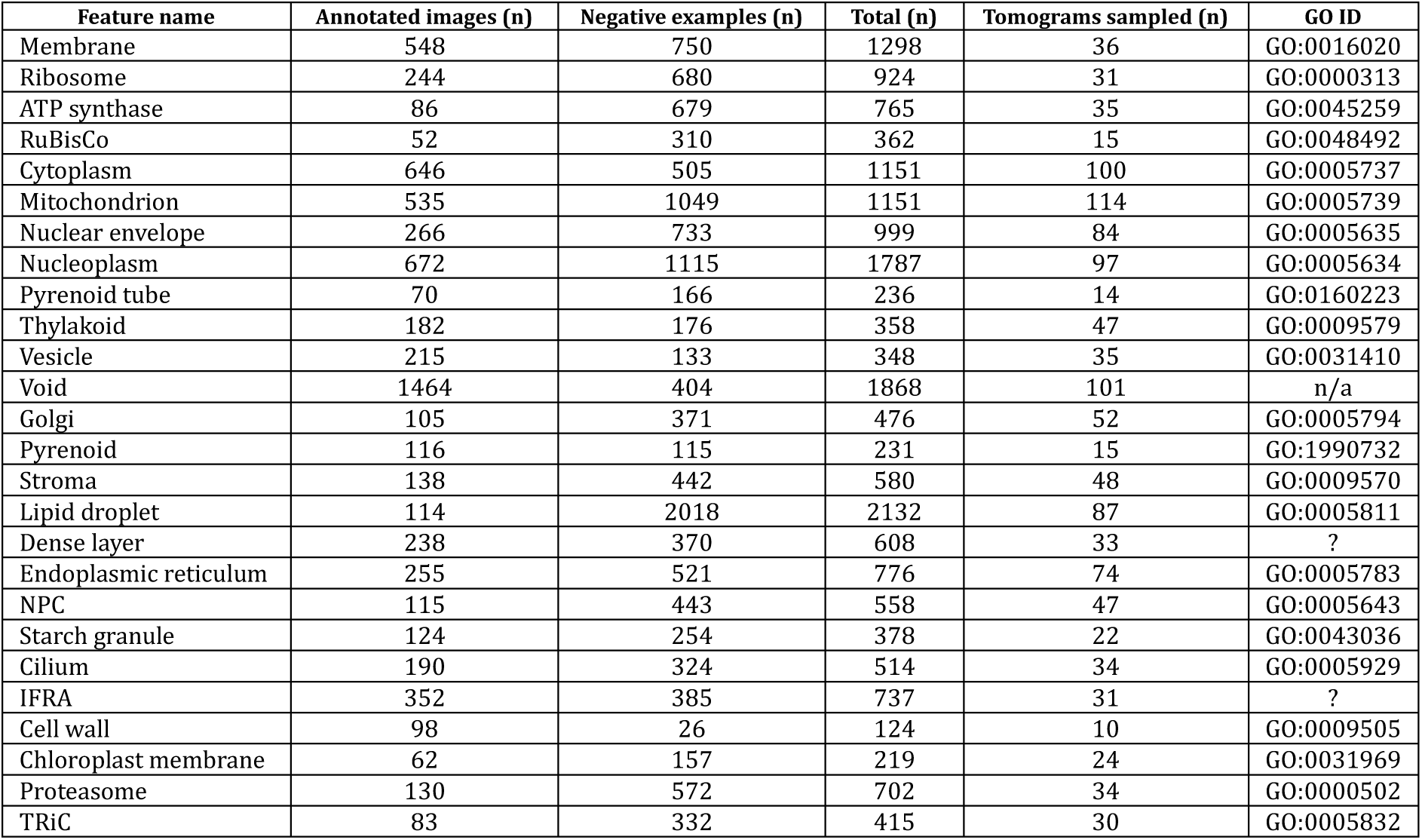
List of segmented features with corresponding Gene Ontology IDs and the sizes of annotated training datasets.

### Network architectures

For the density to macromolecule segmentations we added a new VGGNet-like network architecture^51^ to the Ais network^43^ library. The encoder featured eight repeated 2D convolutional layers with batch normalization, ReLU activation, and 3 × 3 kernel sizes, organised into four blocks with 2 × 2 maximum pooling applied after each block and the number of filters increasing from 128 to 256 to 512 in the last two blocks. The decoder used the same blocks, but with a 2×2 transposed convolutional layer applied after each block and the number of filters decreasing accordingly. The output convolutional layer used a sigmoidal activation function. The models were trained using binary cross-entropy loss and Adam^52^ optimizer with default (tensorflow.keras) settings, for 50 epochs and with a batch size of 32. The trained membrane, ribosome, ATP synthase, and RuBisCo networks are available on the Ais model repository at www.aiscryoet.org.

The architecture of the intermediate network used for the single-feature organelle segmentations was a UNet, similar to the previous model in that it used four blocks in the encoder and decoder with 64, 128, 256, and 512 filters, plus a bottleneck with 1024 filters in between, around which skip connections were used between encoding and decoding convolutional blocks at the same level, in order to better capture details of the spatial context^53^. The joint network was the same, but with double the number of filters in the first encoder and last two decoder blocks. For the intermediary single-feature networks the output layer used a sigmoid activation function, whereas the output of the joint model used softmax activation. The single-feature networks were trained for 100 epochs with batch size 64, using binary cross-entropy as the loss function. The joint model was trained for 300 epochs with batch size 64 using categorical cross-entropy as the loss function and one-hot encoded training labels. Both networks were trained with Adam^52^ as the optimizer with a learning rate of 2.5e-5.

### Data preprocessing

For all training and processing steps, density input images (boxes or full slices) were rescaled to mean 0 and standard deviation 1.0, while macromolecule segmentations were rescaled to a range

-1.0 to +1.0. During training, data was augmented by including all eight permutations of 90° rotations and a single reflection of each image in the input training data. Similar augmentation was applied during inference: volumes were processed four times, in the four 90° rotated orientations, and the resulting segmentations rotated back and averaged.

Immediately after downloading, all tomograms were binned by a factor 2 to a size of 512 × 512 × 256 voxels in order to save both disk space and processing time, while also retaining sufficient resolution to segment the smallest features (ATP synthase, RuBisCo, membranes). Macromolecule segmentations were performed using these volumes (7.84 A /pixel). The volumes were binned by another factor of 2 to a size of 256 × 256 × 128 voxels (15.68 A /pixel) prior to computing the organelle segmentations.

### Data visualization

All images in this article were rendered with Ais and Pom, the Python module that was created during this work. Pom contains features for automated rendering of 3D scenes with user-defined compositions. By default, the command ‘pom render’ outputs two images for each tomogram, featuring: i) all macromolecules, and ii) the top-3 organelles with the largest volume fraction in that tomogram. Alternative configurations, such as cytoplasm + nucleoplasm + nuclear envelope + membrane (e.g. Figure 2 and Figure 2B supplement 3) are also supported; volumes can be rendered either as isosurfaces (e.g. Figure 2A) or emissive volumes (e.g. Figure 2B). Projection images were generated with the command ‘pom projections’. Rendering all the 3658 3D images and 76818 2D projection images that are used in the dataset summary (cryopom.streamlit.app) took around 30 minutes (64 concurrent processes on one Linux server).

### Template matching

Our OpenGL-based implementation of GPU-accelerated real-space 2D template matching was fairly rudimentary and not optimized beyond what we needed to perform the tests outlined above. In brief, the steps employed in these processes were as follows: first, we resampled an N × N × N-sized template density volume and mask volume each in K different orientations, as specified by a list of transform matrices, and uploaded a serialized list of the central slice of each resampled template T_i_ and mask M_i_ to the GPU as an OpenGL shared storage buffer object (SSBO). Next, for every slice in a tomogram volume, we uploaded the slice and the corresponding area-selective mask to the GPU as grayscale 32-bit float textures. Next, we dispatched one compute shader work group for every pixel in the image simultaneously, with each work group comprising K shader invocations. GPU-side, the 0^th^ invocation first copied image I, of size N × N and centred on its group’s pixel coordinate, into shared local memory. Next, each invocation computed the Roseman local correlation coefficient for images I and T_i_, scoring voxels specified by mask M_i_ only, and store it in local memory as a score S_i_. Finally, invocation 0 would find the maximum score S and write both S and the corresponding index i into an output texture, which is downloaded to the CPU. For ASTM of NPCs we used a template with a box size of 64, resampled in 500 orientations that were uniformly distributed on a unit sphere between polar angles -40 to +40°, with no further rotation applied around this orientation vector.

For ASTM of the lightbulb complexes, masks were generated by thresholding mitochondrion segmentations (>0.5) and using 2D binary erosion and dilation to generate a shell of ∼20 nm thickness around the outside of mitochondria. We also generated void and thylakoid masks (thresholding at >0.2) and subtracted these from the mitochondrion masks, to avoid performing TM in regions of low quality or where segmentations were somewhat ambiguous. The 32 × 32 × 32 template was picked manually and blurred (20 A st. dev. Gaussian) and a template mask generated by annotating the template in Ais^43^; both were resampled in 500 orientations spaced approximately uniformly on a unit sphere between polar angles of -30° and +30° and then reduced to 2D as before. Input density volumes were 512 × 512 × 256 pixels, but to strike a balance between processing speed and accuracy we applied a stride of 2 to the mask, meaning every 2^nd^ voxel (along all three axes) of the volume mask was set to 0 (resulting in 8-fold faster processing). Because the blurring kernel applied to the template was larger than one pixel, we assumed that applying this stride would not significantly reduce accuracy.

### Particle picking

To pick particles, we used essentially the same procedure as is implemented in Ais^43^, but adapted the code so that it could be run in parallel on multiple CPUs. The process involved thresholding the ribosome segmentations, computing a distance transform followed by a watershed transformation, and then considering the maxima in the resulting transformed volume to be coordinates that corresponded to ribosome positions. The accuracy this approach depends on the accuracy of the network used for segmentation and on a number of tuneable parameters (threshold level, minimum particle size, minimum particle spacing). Particle sets resulting from automated picking based on neural network segmentations typically require further curation, e.g. by classification, in order to discard false particles; the number of picked ribosomes (roughly 300,000) is thus likely an overestimate of the actual number present in the data.

## Software availability

A command-line interface tool for the training and processing steps discussed in this article is available on the Python package index as *‘Pom-cryoET’*, with source code at github.com/bionanopatterning/Pom and documentation at pom-cryoet.readthedocs.io. The code for the template matching experiments is available via github.com/bionanopatterning/Pommie. A new version of Ais, with added features that help organise the large number of annotations and significantly improved processing speed, is available at github.com/bionanopatterning/Ais and on PyPi as ‘*Ais-cryoET’* (version 1.1.40 or higher). All are available under the GNU GPLv3 open source license.

## Data availability

The original data by Kelley and Khavnekar et al. was downloaded from the CZII CryoET Data Portal, available via cryoetdataportal.czscience.com/datasets/10302. All segmentation volumes are also available on the portal under accession number 10314. We are in the process of uploading all segmentation volumes to the portal as well, but these data are currently not yet available. An automatically generated segmentation report is hosted at cryopom.streamlit.app. This example includes projection images of all segmentations, 3D visualizations of all macromolecule segmentations and top 3 organelle components for each tomogram, and an interactive summary table.

## Acknowledgements

This work was supported by the following grants to THS: European Union’s Horizon Europe Program IMAGINE grant 101094250, and the Netherlands Organization for Scientific Research Grant VI.Vidi.193.014.

This work would have not been possible without the free availability of a number software packages including Python, glfw^54^, PyOpenGL^55^, streamlit^56^, tensorflow^57^, keras^58^, mrcfile^59^, numpy^60^, scipy^61^, scikit-image^62^, and imgui^63^.

We thank Leoni Abendstein, Frank Faas, Bram Koster, Montse Barcena, and Meindert Lamers for valuable discussions and feedback. We are especially grateful also to the authors of the 1829 tomogram *C. reinhardtii* dataset for making these data freely available online. Finally, we thank the participants of the 2024 Gordon Research Conference on 3DEM, where the idea for this project was born, and Utz Ermel and colleagues at the Chan Zuckerberg Imaging Institute for their help in making our and many other cryoET datasets and segmentations available online..

## Supplementary Figures

**Figure 1 supplement 1.**
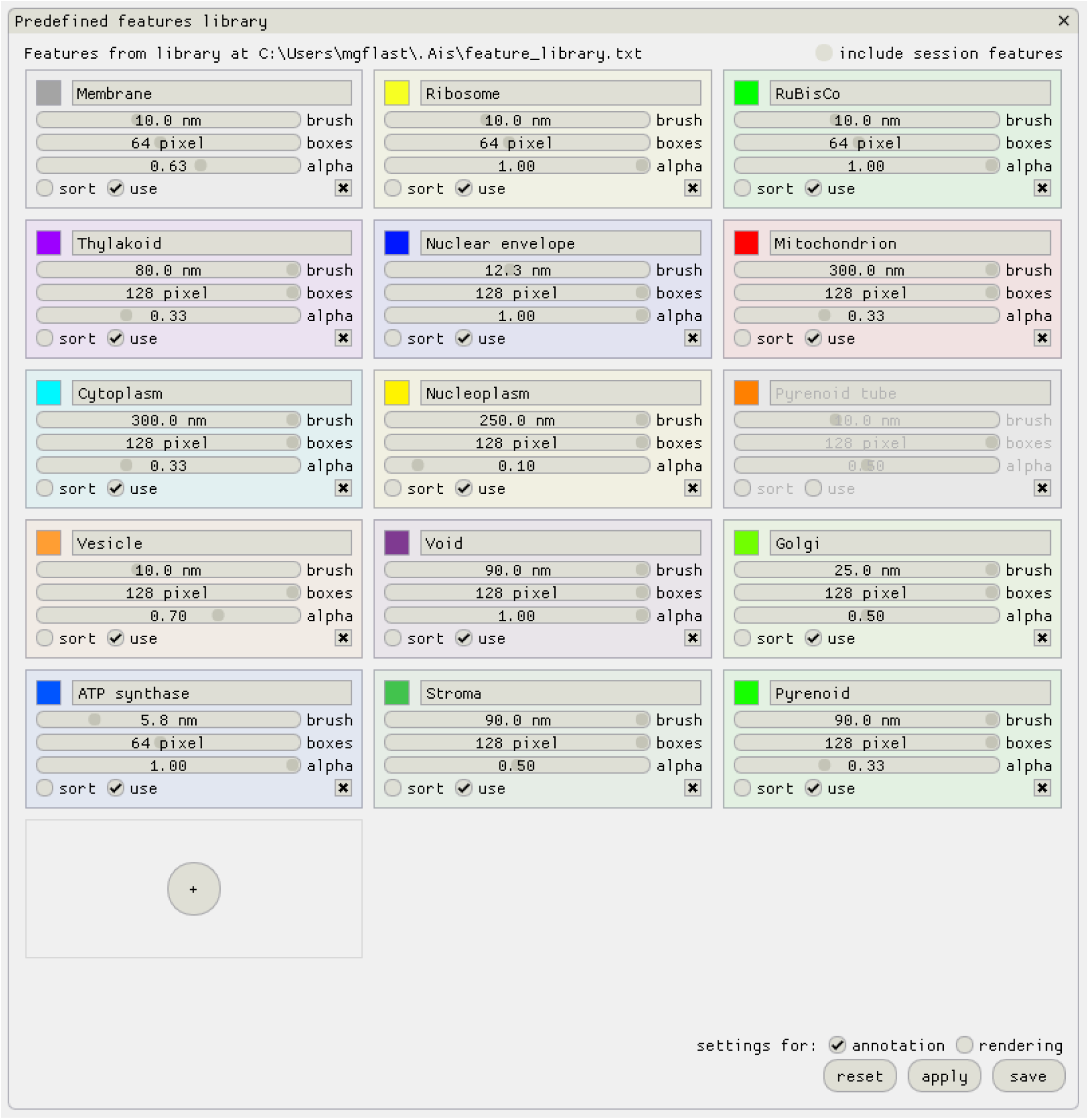
The Ais ‘feature library’. During this project we realised that additional tools for organising a large number of annotations would be helpful. We added a feature library to Ais, allowing users to configure named presets for annotation (e.g. ‘Membrane’), and choose default colours and visualization settings. During annotation and rendering, the same colours and parameters are than always automatically used for the same feature. When rendering 3D visualizations with Pom (the Python module accompanying this article) the visualization settings defined in the Ais feature library are also automatically used.

**Figure 2 supplement 1.**
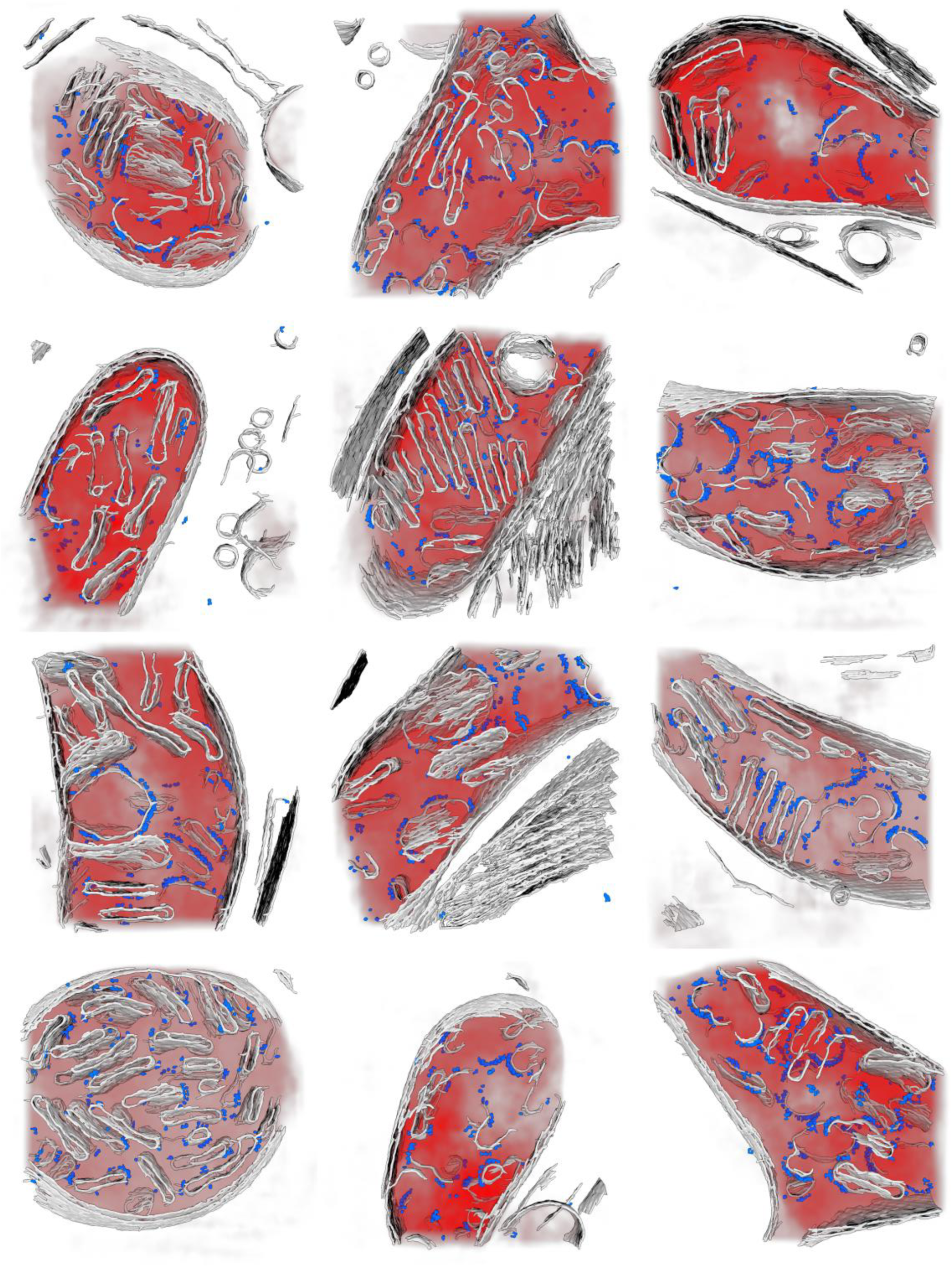
An exemplary sub-dataset: tomograms containing mitochondria. An automatically selected subset of tomograms containing mitochondria (filtering for >20% mitochondria content) rendered in 3D. The visualization shows membrane (gray), ATP synthase (blue), and mitochondrion (red) segmentations, with macromolecules rendered as isosurfaces and mitochondria as an emissive volume.

**Figure 2 supplement 2.**
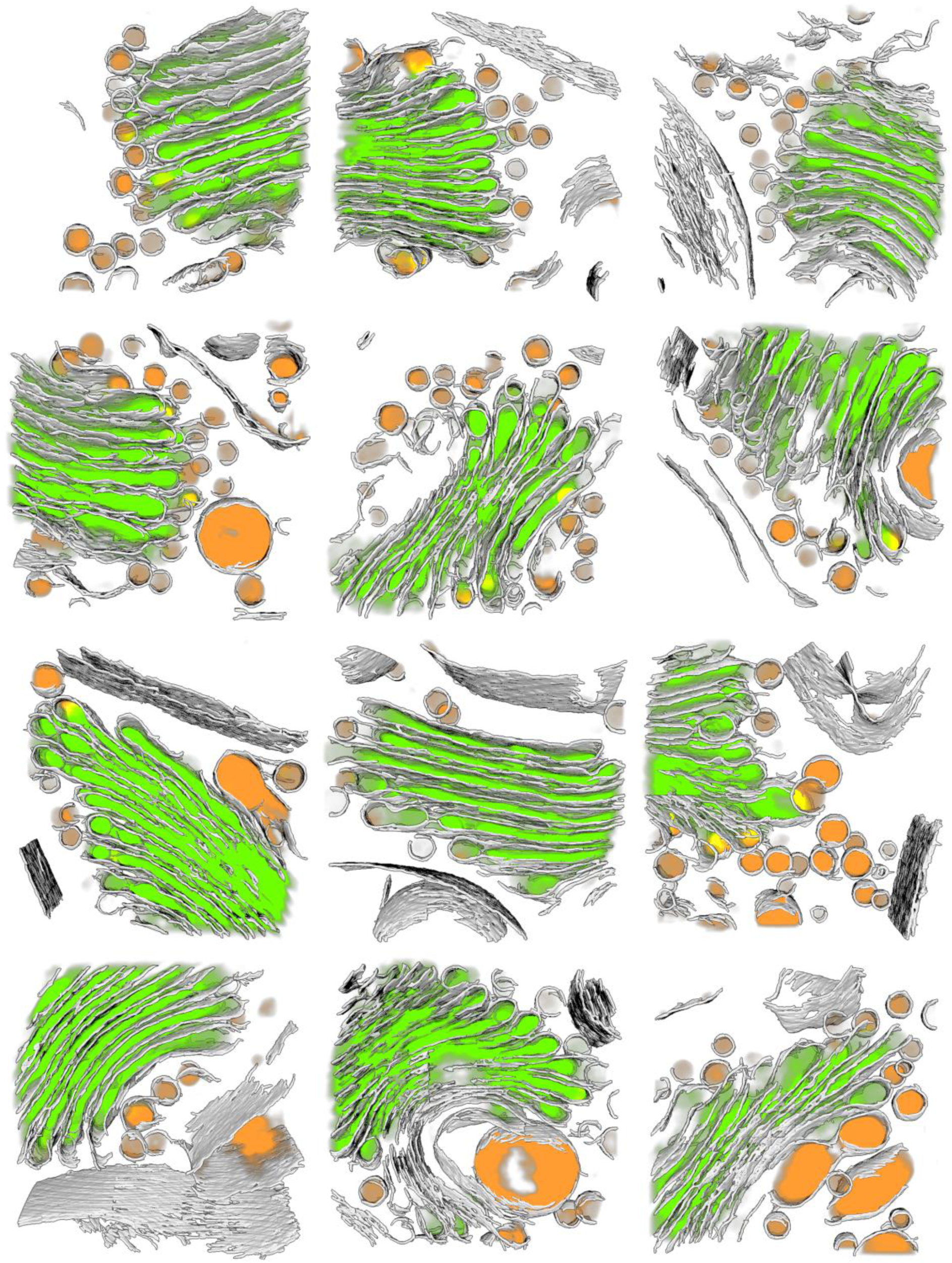
An exemplary sub-dataset: tomograms containing the Golgi complex. An automatically selected subset of volumes containing Golgi (filtering for >10% Golgi content) rendered in 3D. The visualization shows membrane (gray), vesicle (orange), and Golgi (green) segmentations, with membranes rendered as isosurfaces and vesicle and Golgi segmentations as emissive volumes.

**Figure 2 supplement 3.**
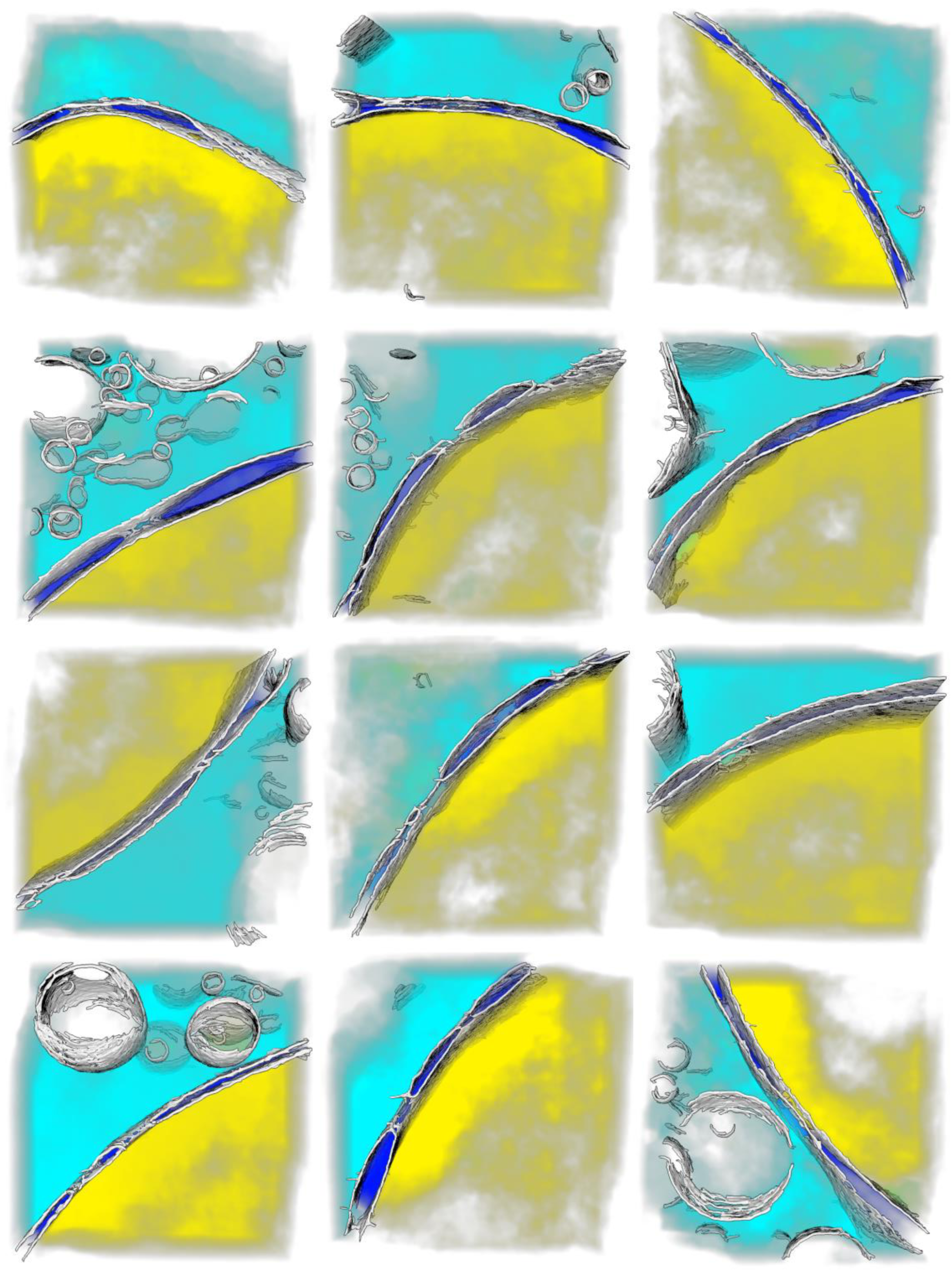
An exemplary sub-dataset: volumes that contain both nucleoplasm and cytoplasm. A number of automatically selected volumes containing both nucleoplasm and cytoplasm (filtering for >10% of both) rendered in 3D. The visualization shows membrane (gray), nucleoplasm (yellow), cytoplasm (cyan), and nuclear envelope (blue) segmentations, with membranes rendered as isosurfaces and the rest as emissive volumes.

**Figure 2 supplement 4.**
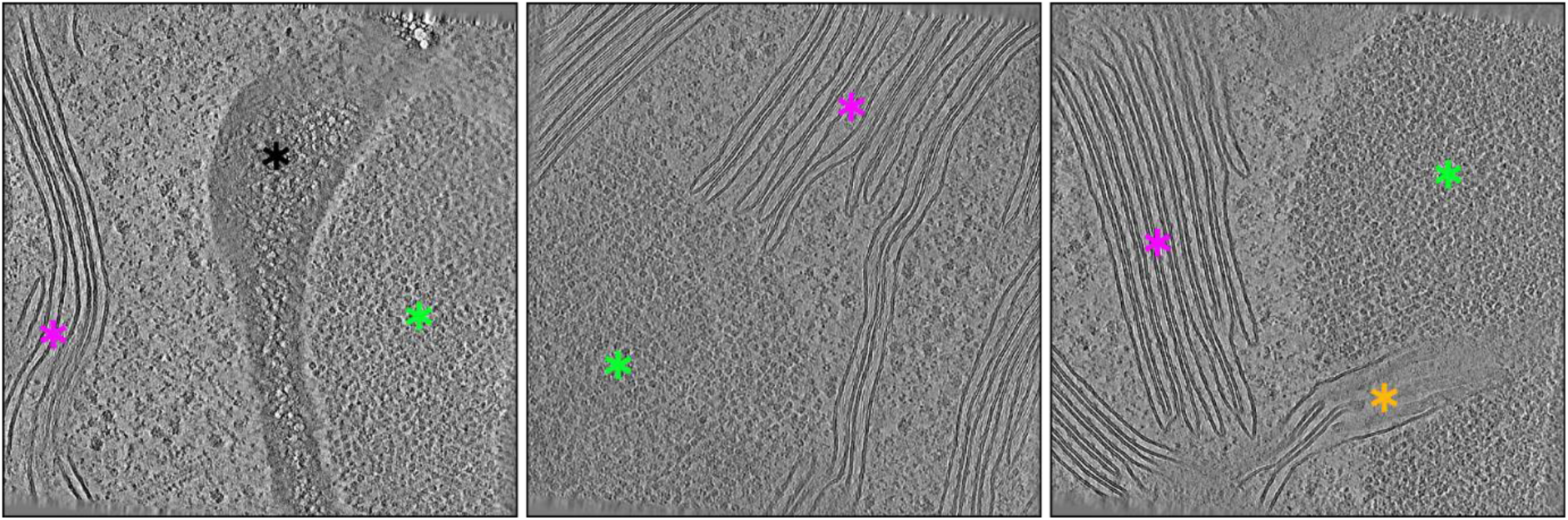
A number of exemplary tomograms with a rare composition, featuring both thylakoid membranes and the pyrenoid. Although both thylakoid membranes (magenta) and pyrenoids (green) are common in the dataset (n = 651 for tomograms containing at least 5% thylakoid, n = 74 for at least 5% pyrenoid), instances where both features are visible within the same tomogram are rare (n = 7 for both thylakoid and pyrenoid >5%). By specifically selecting these tomograms a number of interesting structures can be found, such as a section of the starch sheath that envelops the pyrenoid (black, left panel), thylakoid membranes (magenta) and pyrenoid (green) in close proximity (center), and a view of thylakoid membranes branching off into a pyrenoid, forming a pyrenoid tube (orange, right panel).

**Figure 3 supplement 1.**
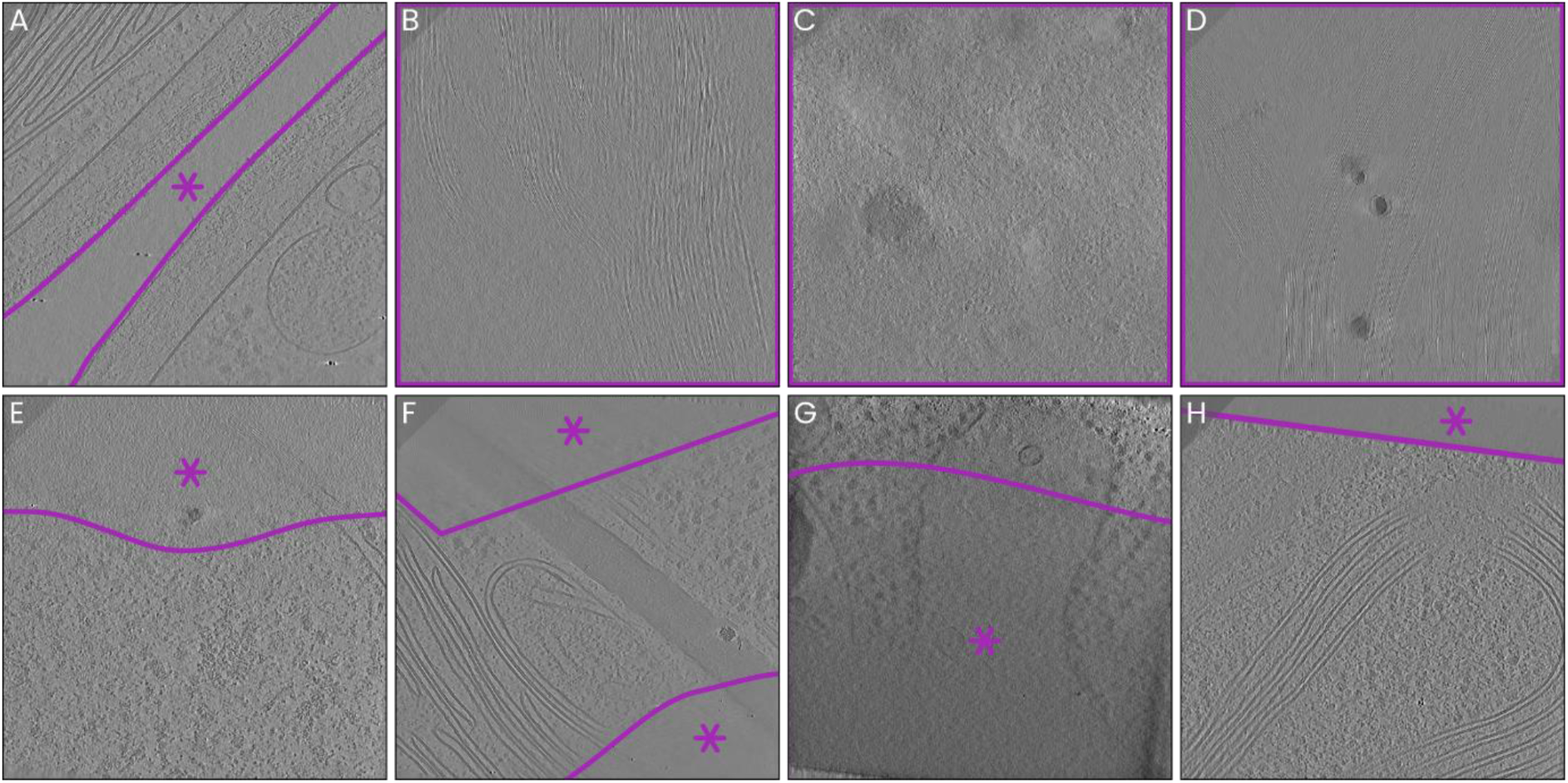
Examples of image features considered void. Areas considered void (marked with magenta asterisks and borders) included those not containing cellular material (**A**), slices or sections of slices that lie above or below the edges of the lamella (**B-F**), areas with low image quality (**G**), and reconstruction artefacts (**F**).

**Figure 4 supplement 1.**
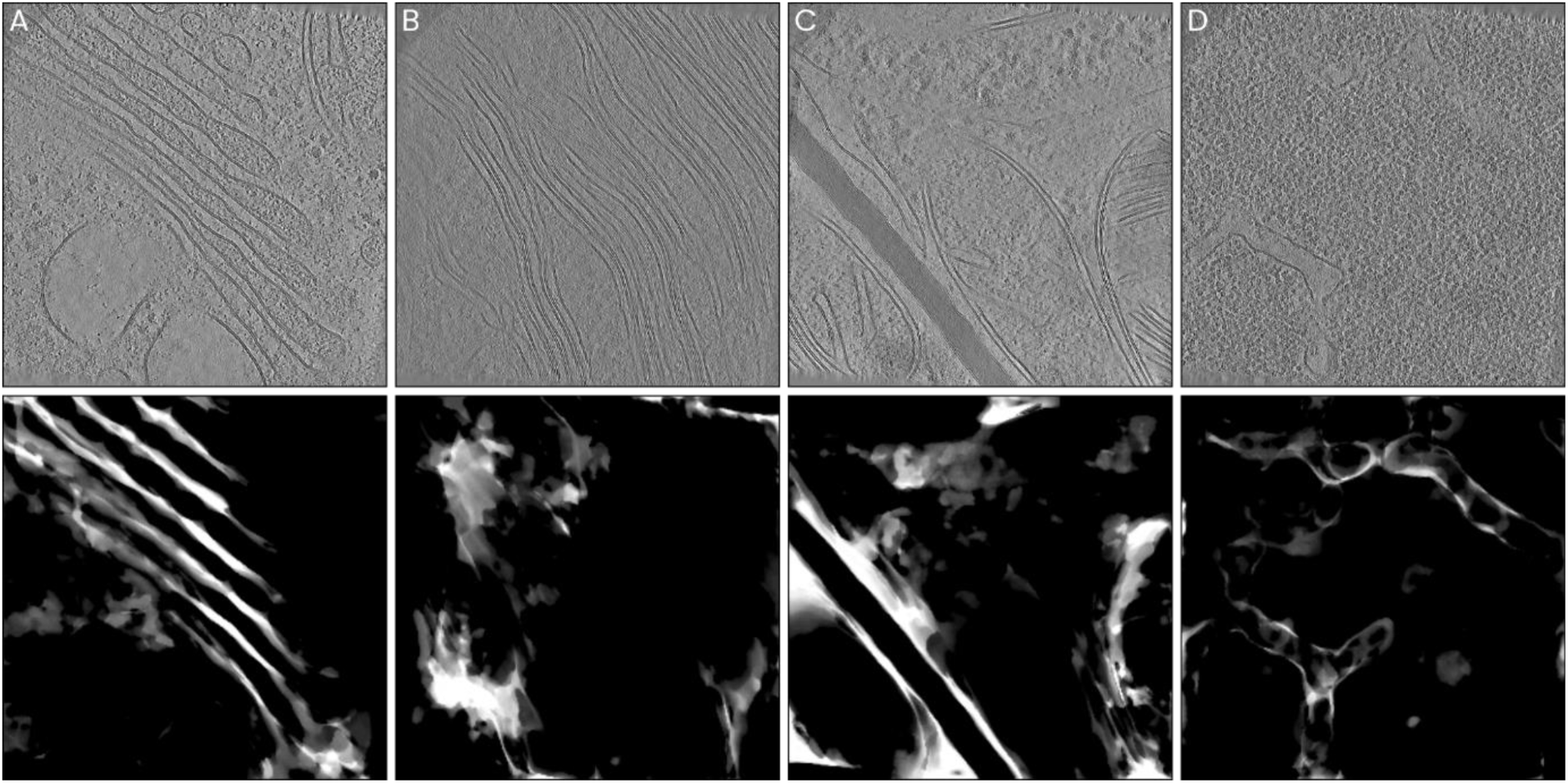
Examples of ‘artefactual’ unknown segmentation outputs. The top row shows central slices of tomograms, while the bottom row shows z-projections each corresponding unknown-segmentation volume. **A)** Regions of cytoplasm located in between Golgi stacks were not included as cytoplasm-annotated images in the cytoplasm training dataset, and are segmented as unknown as a result. **B)** Regions within chloroplasts where the orientation of the membranes led is such that differentiating between stroma and thylakoid is difficult, resulting in unknown annotations instead. **C)** Narrow regions of cytoplasm in between other structures are poorly recognized and annotated as unknown instead. **D)** Interfaces between two classes, in this case between pyrenoid tubes and pyrenoid, are also sometimes segmented as unknown. This effect can occur when the boundary between two features (such as a membrane) is not consistently included in the training annotations for at least one of the two features. Some of these artefacts could perhaps be avoided by applying post-processing filters to the unknown segmentations.

**Figure 4 supplement 2.**
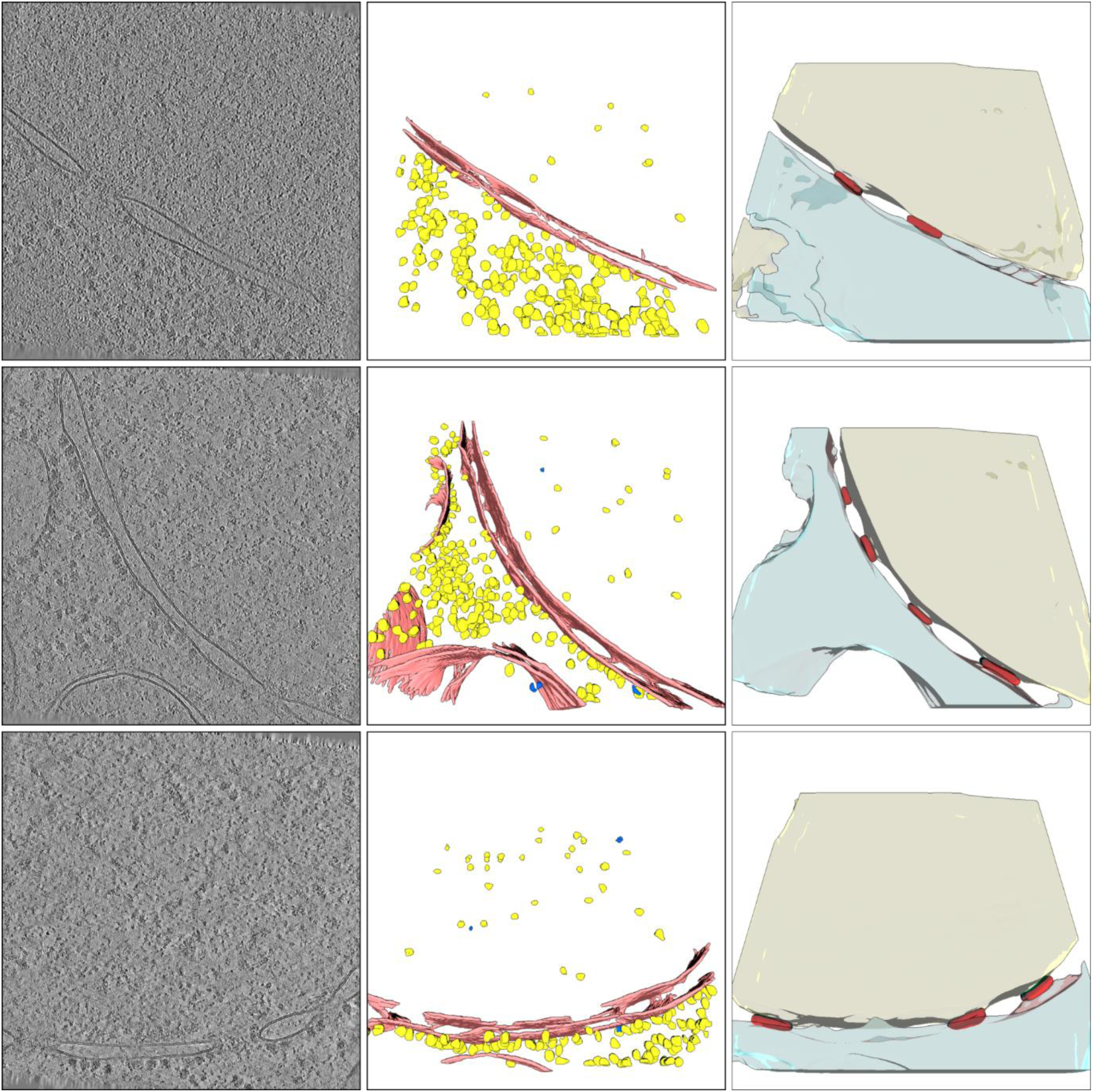
Examples of nuclear pore complex (NPC) segmentation results after expanding the ontology to include previously unknown components. Left column shows density (central slice of tomograms), middle shows macromolecule segmentations (membranes in red, ribosomes in yellow, ATP synthase in blue). Note that some of the ATP synthase particles in these examples are probably false segmentations. Right shows organelle segmentations (nucleoplasm in beige, NPC in red, cytoplasm in cyan).

**Figure 4 supplement 3.**
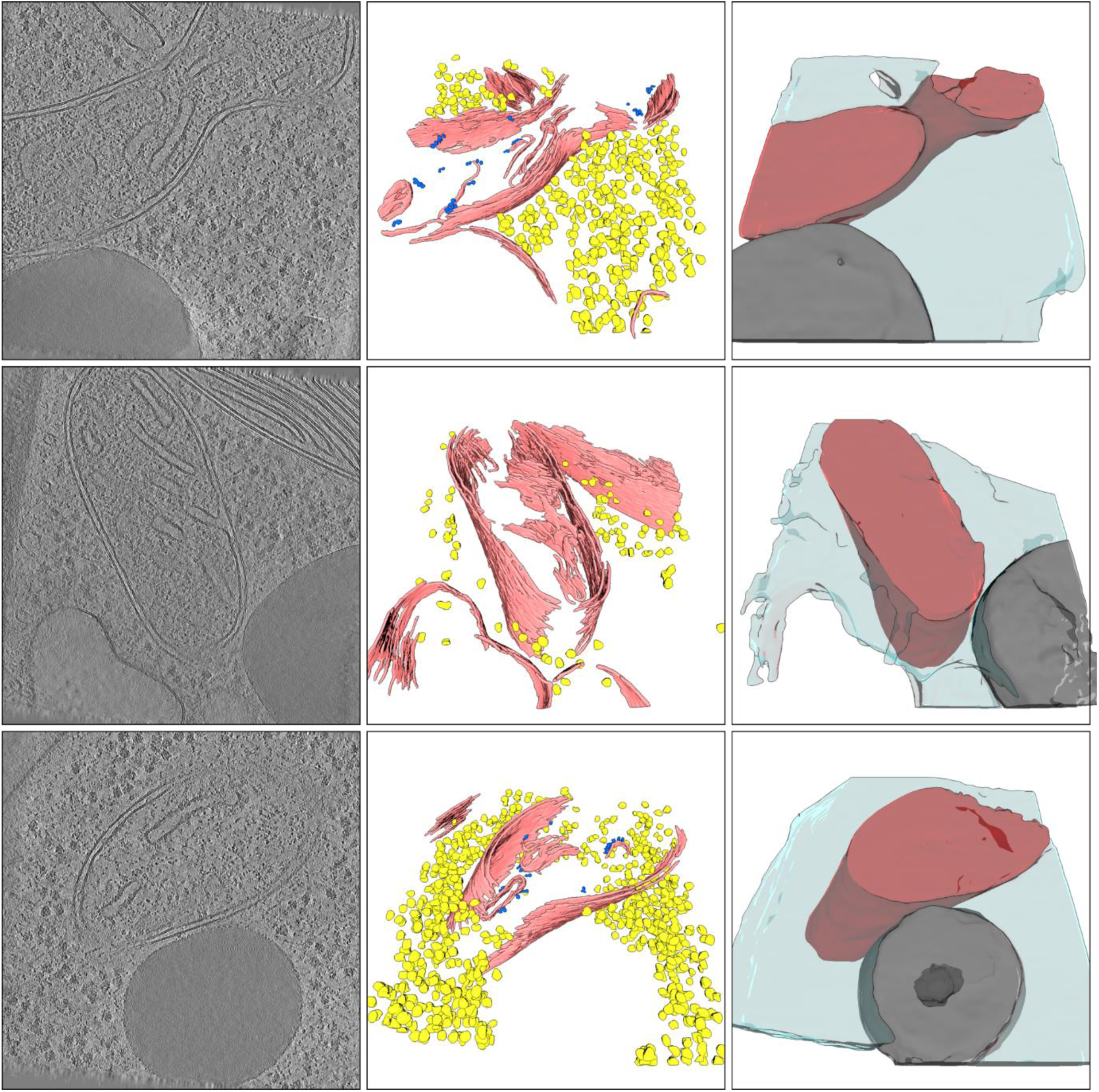
Examples of lipid droplet (LD) segmentation results after expanding the ontology to include previously unknown components. Left column shows density (central slice of tomograms), middle shows macromolecule segmentations (membranes in red, ribosomes in yellow, ATP synthase in blue), right shows organelle segmentations (cytoplasm in cyan, mitochondria in red, lipid droplets in dark gray).

**Figure 4 supplement 4.**
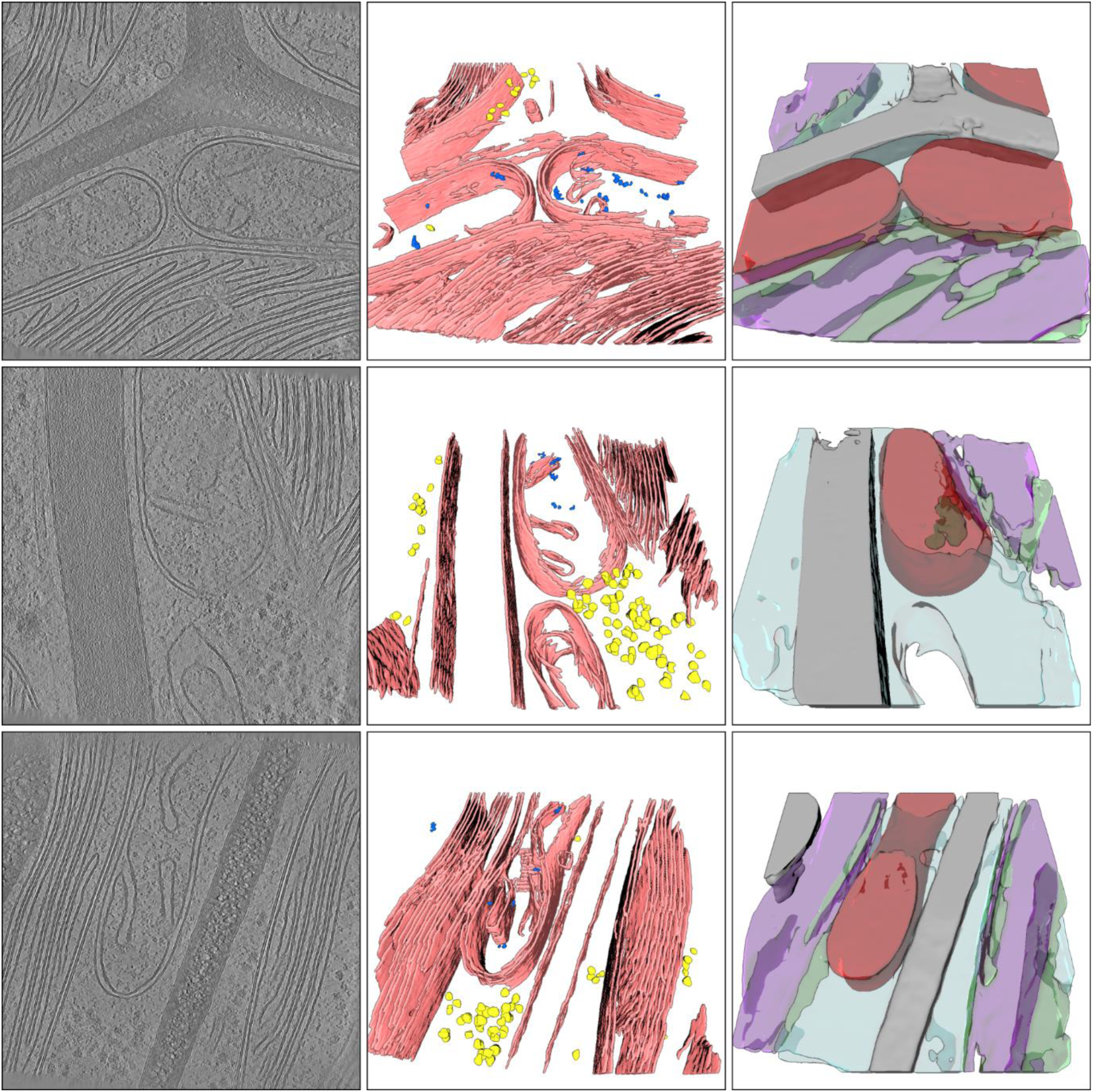
Examples of dense layer (DL) segmentation results after expanding the ontology to include previously unknown components. Left column shows density (central slice of tomograms), middle shows macromolecule segmentations (membranes in red, ribosomes in yellow, ATP synthase in blue.), right shows organelle segmentations (cytoplasm in cyan, mitochondria in red, thylakoid in purple, stroma in green, the dense layer in gray).

**Figure 4 supplement 5.**
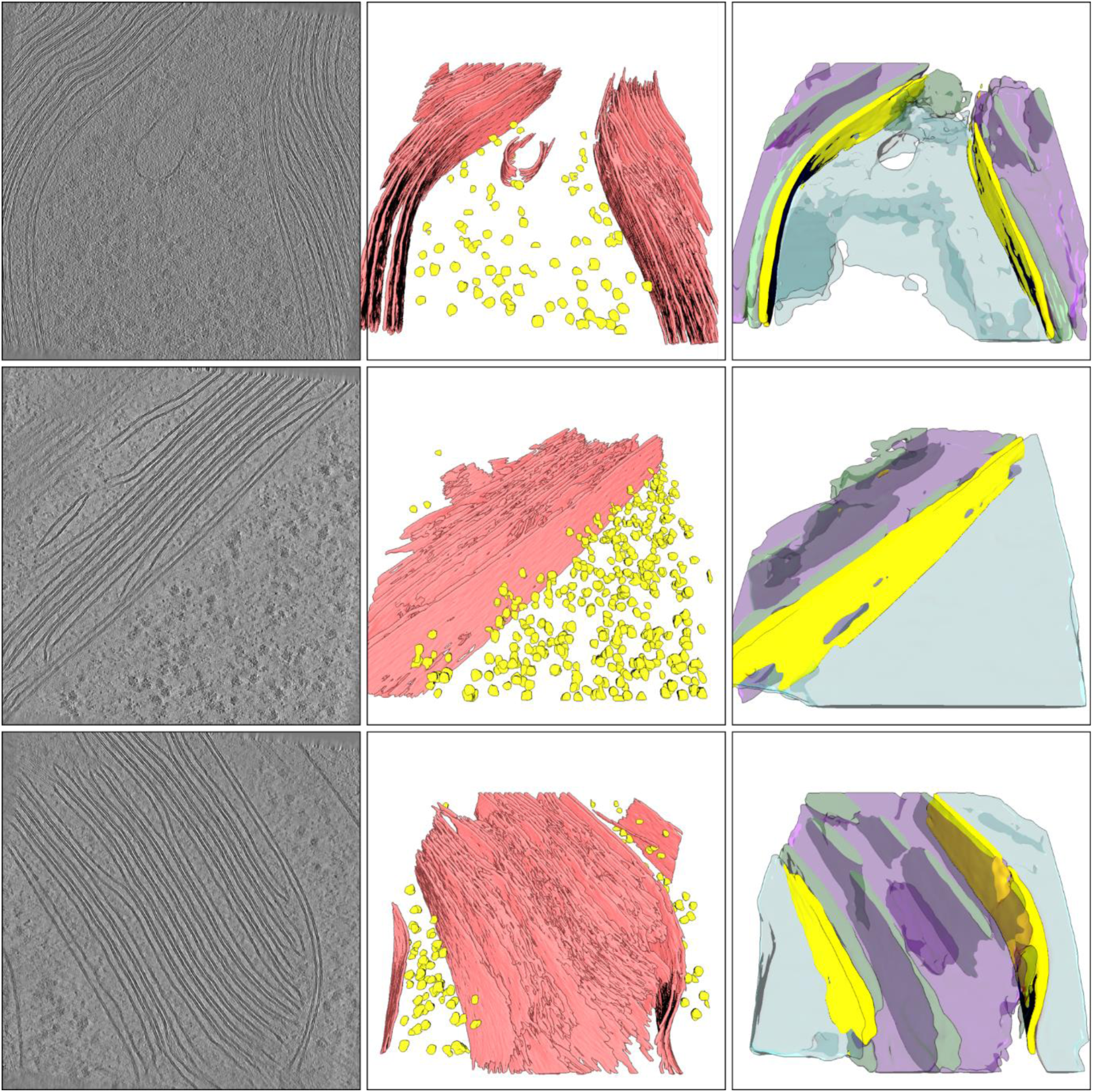
Examples of chloroplast outer membrane segmentation results after expanding the ontology to include previously unknown components. Left column shows density (central slice of tomograms), middle shows macromolecule segmentations (membranes in red, ribosomes in yellow), right shows organelle segmentations (cytoplasm in cyan, thylakoid in purple, stroma in green, chloroplast outer membrane in yellow).

**Figure 4 supplement 6.**
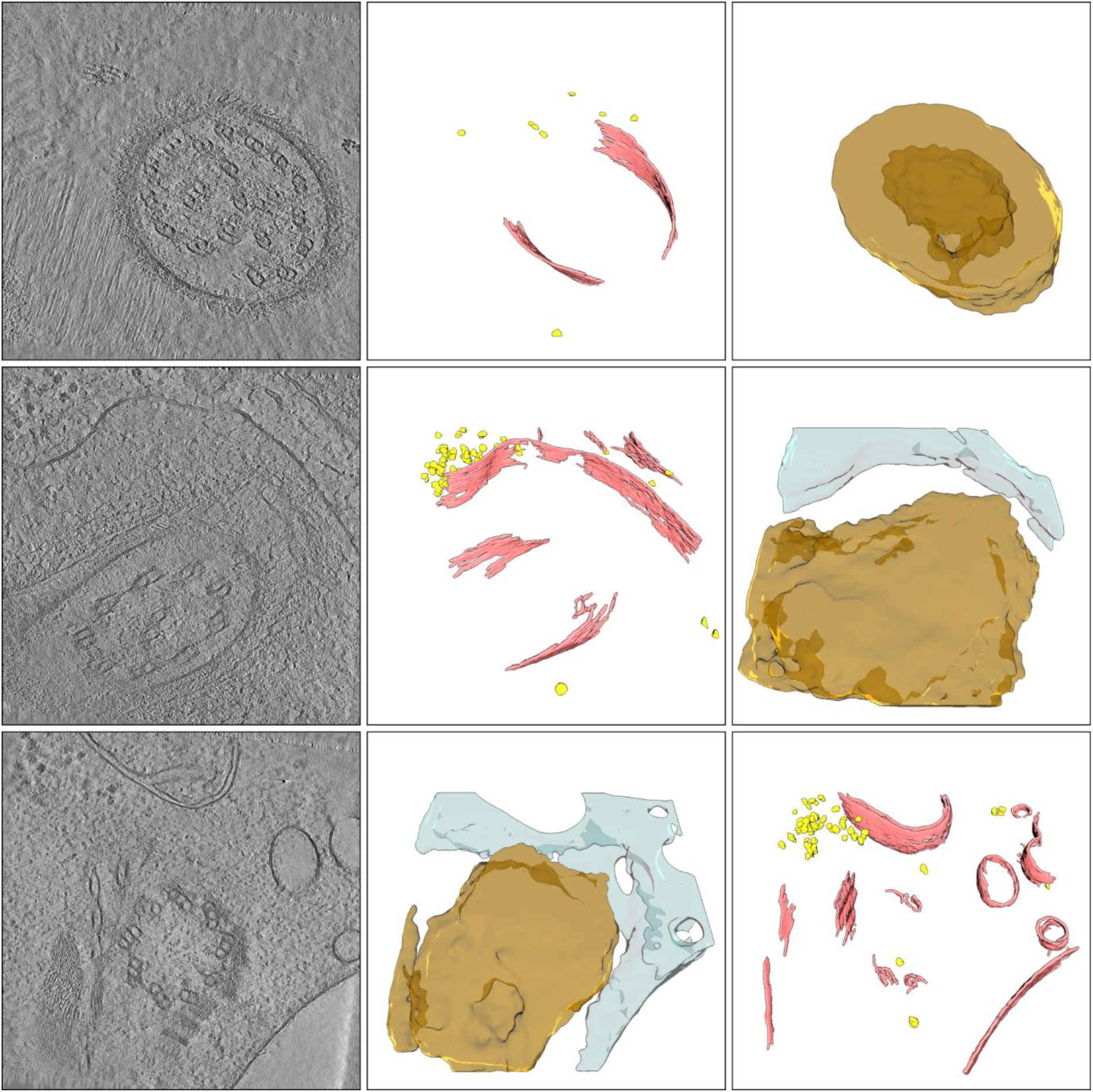
Examples of cilium segmentations after expanding the ontology to include previously unknown components. Left column shows density (central slice of tomograms), middle shows macromolecule segmentations (membranes in red, ribosomes in yellow), right shows organelle segmentations (cytoplasm in cyan, cilium in orange). Note that the cilum annotations and resulting segmentations were not very precise; we roughly annotated the microtubule bundles, surrounding membrane and glycocalyx, and, when visible, also parts of the interflagellar transport train and nearby microtubules.

**Figure 4 supplement 7.**
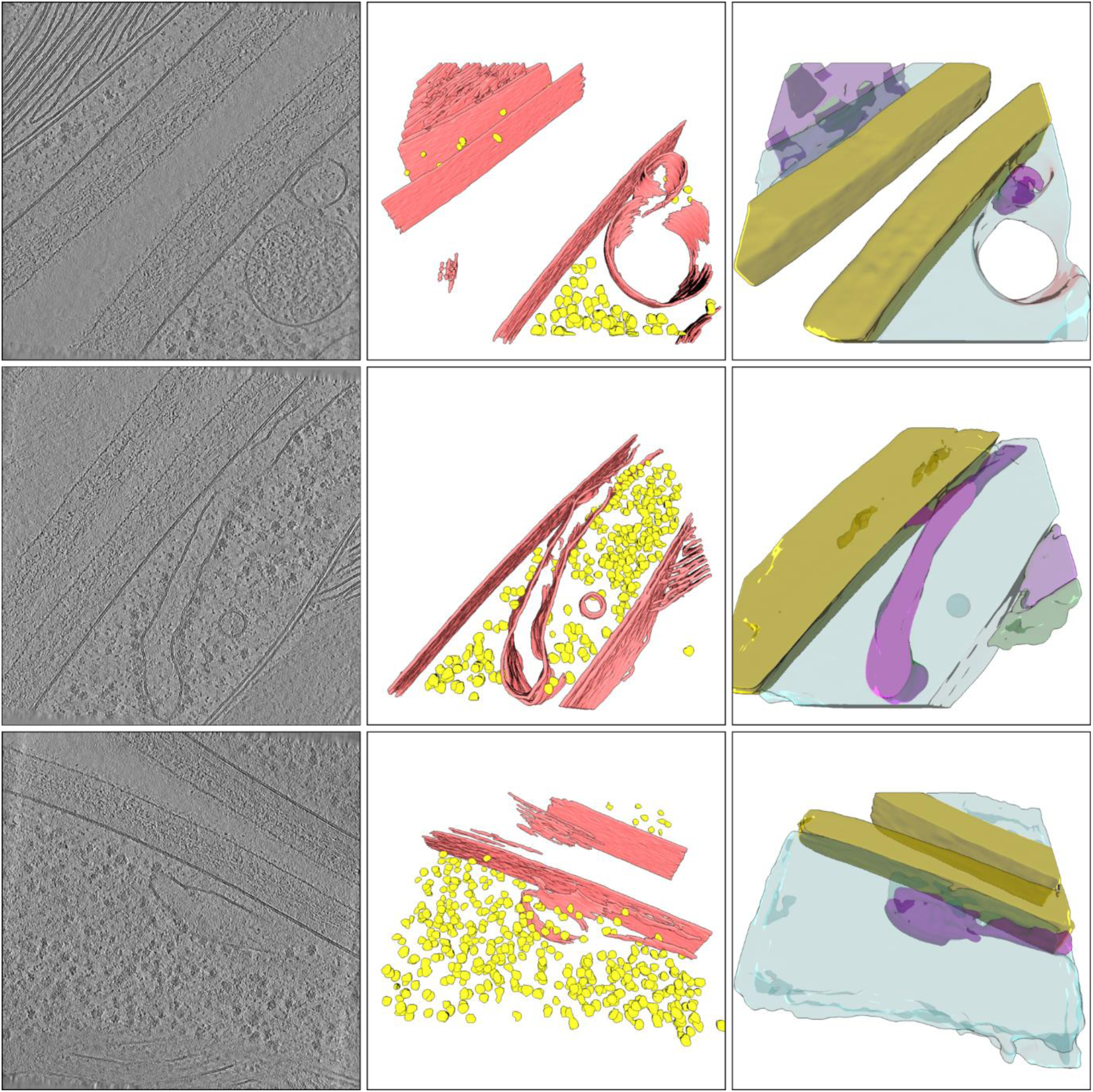
Examples of cell wall segmentations after expanding the ontology to include previously unknown components. Left column shows density (central slice of tomograms), middle shows macromolecule segmentations (membranes in red, ribosomes in yellow), right shows organelle segmentations (cell wall in ochre, cytoplasm in cyan, thylakoid (top row only) in violet, stroma in olive, endoplasmic reticulum in magenta).

**Figure 4 supplement 8.**
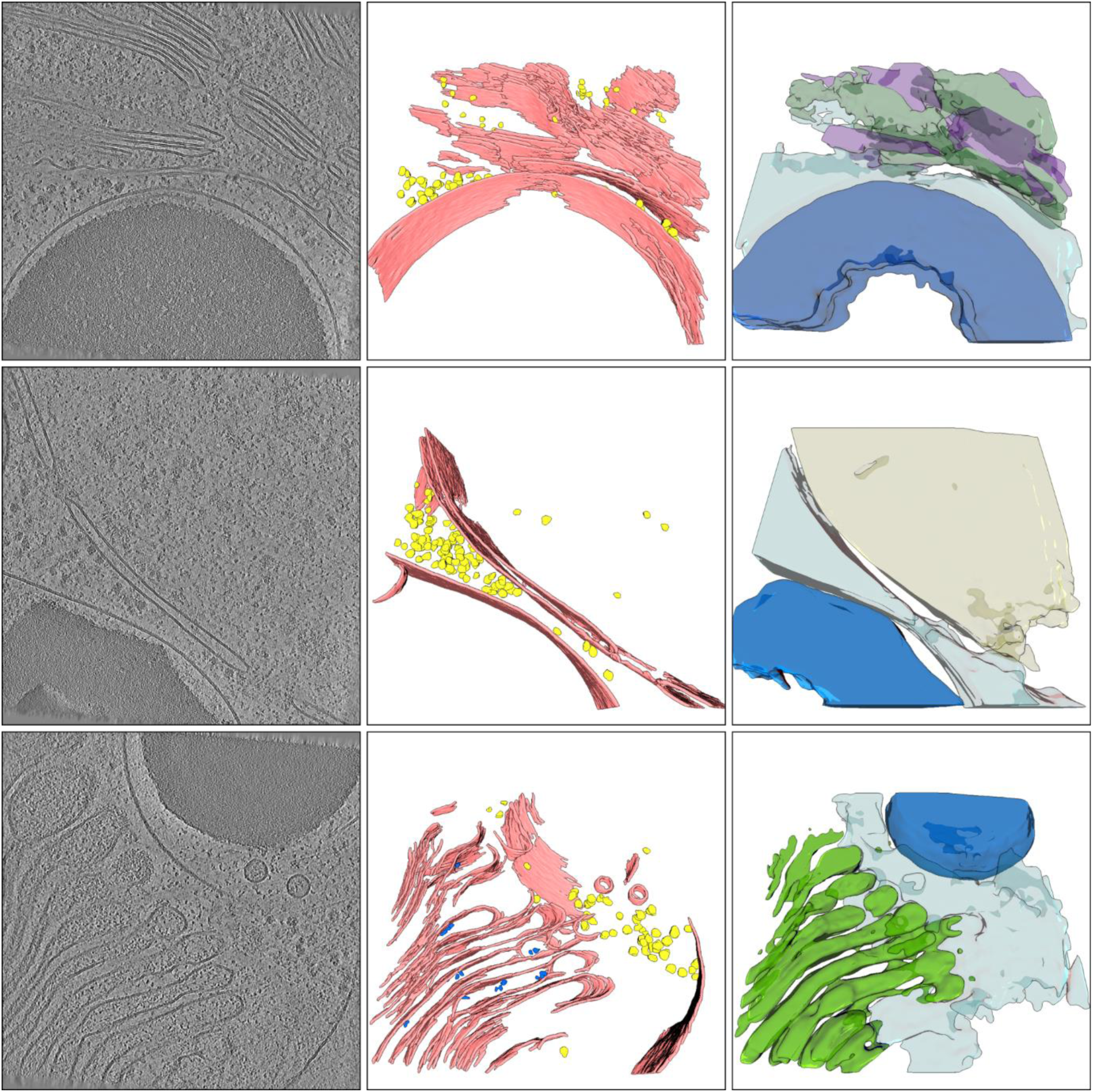
Examples of starch granule segmentations after expanding the ontology to include previously unknown components. Left column shows density (central slice of tomograms), middle shows macromolecule segmentations (membranes in red, ribosomes in yellow, ATP synthase in blue). Note that the ATP synthase particles in the bottom row are likely false detections. Right shows organelle segmentations (starch granule in blue, cytoplasm in cyan, thylakoid in purple, stroma in olive, nucleoplasm in beige, Golgi in green).

**Figure 4 supplement 9.**
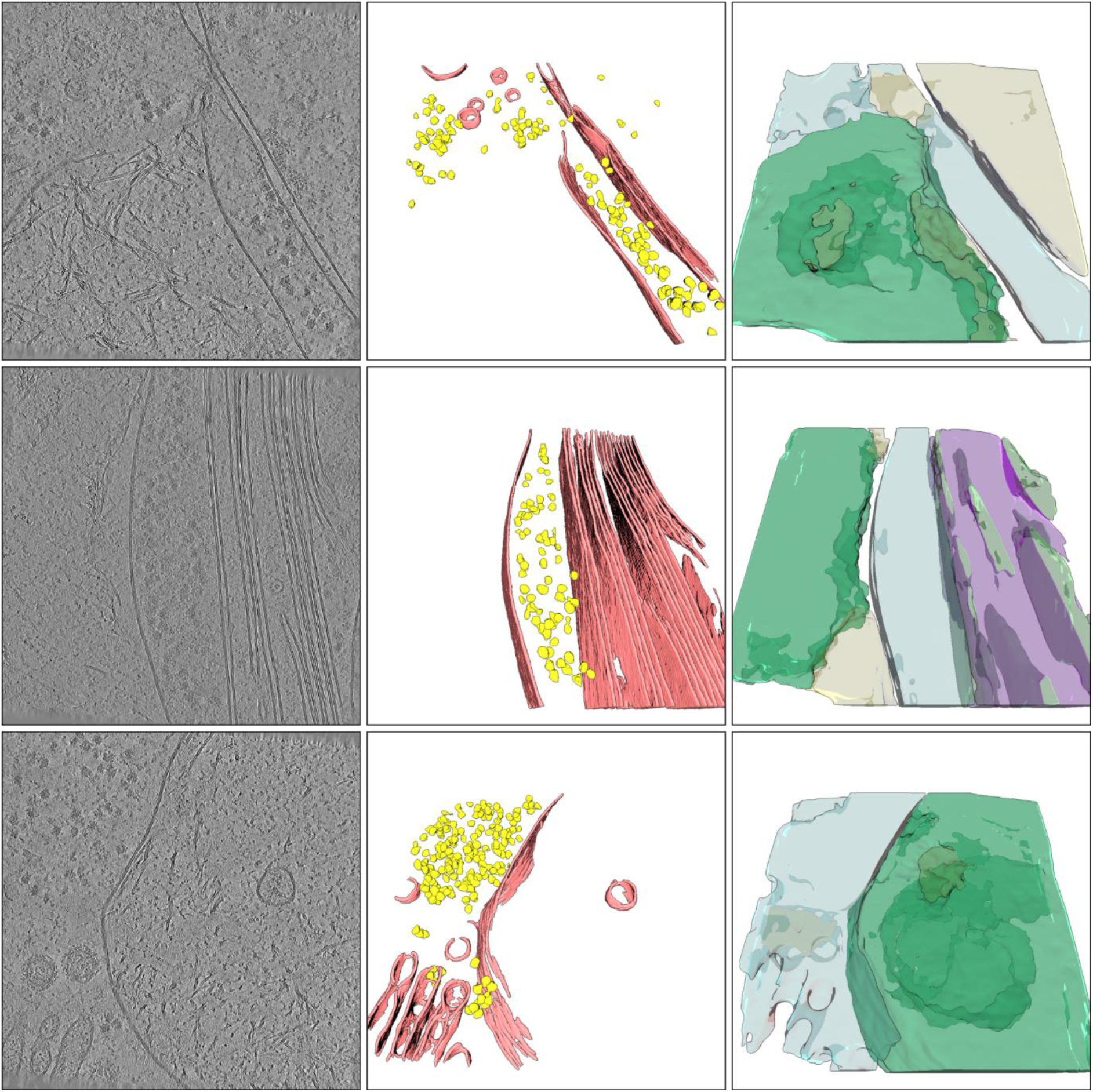
Examples of ‘intermediate-filament-rich areas’ (IFRA) segmentations after expanding the ontology to include previously unknown components. Left column shows density (central slice of tomograms), middle shows macromolecule segmentations (membranes in red, ribosomes in yellow). Note that intermediate filaments themselves were not segmented (attempts to do so led to many sections of membrane falsely being annotated as intermediate filaments instead, so we decided not to include this macromolecular feature for the time being). Right shows organelle segmentations (IFRA in green, cytoplasm in cyan, nucleoplasm in beige, thylakoid in purple, stroma in olive).

**Figure 5 supplement 1.**
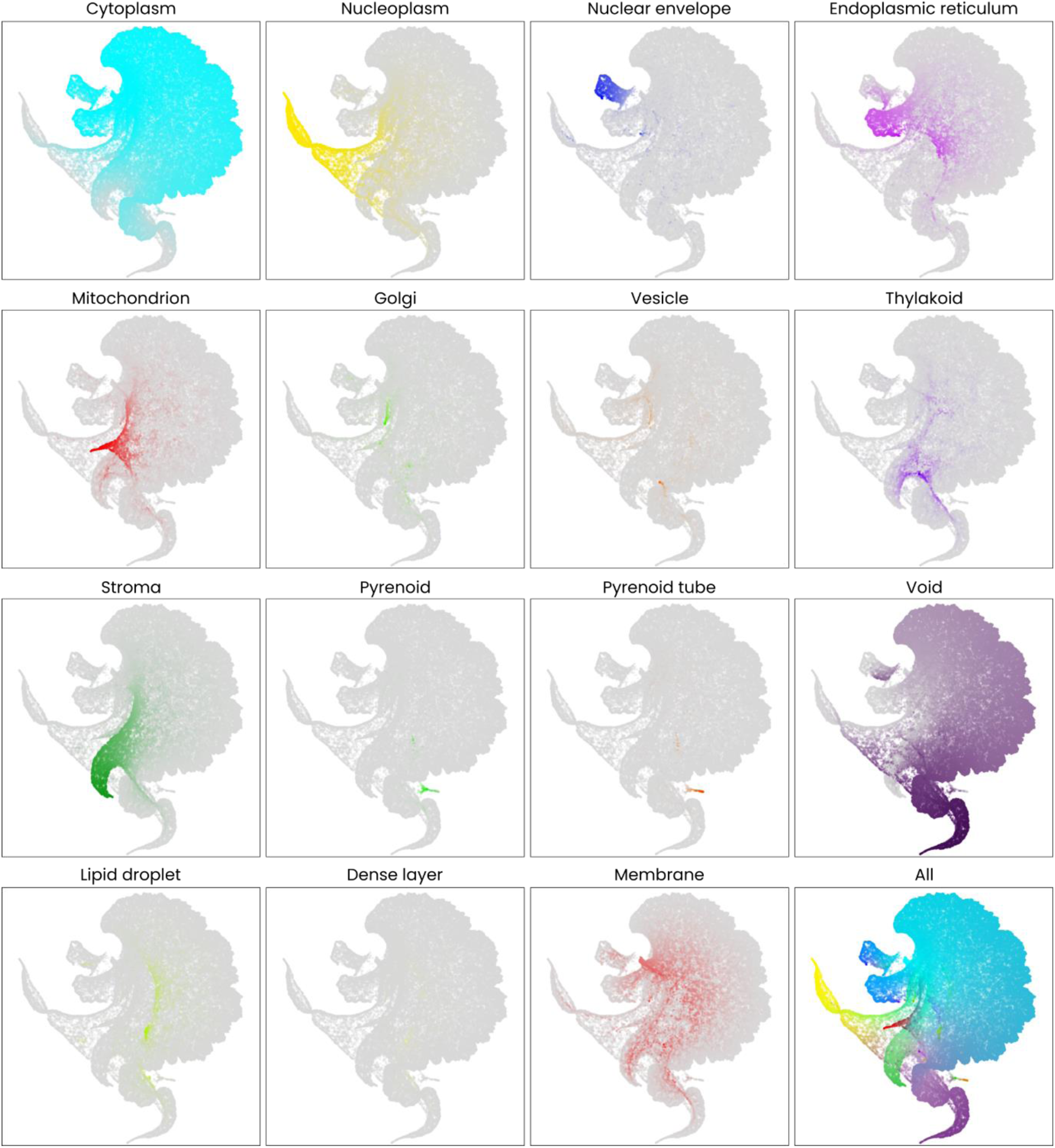
UMAP of ribosome context vectors with specific contextual features highlighted. Aside from cytoplasmic, nuclear, endoplasmic reticulum associated and nuclear envelope associated ribosomes, ribosomes (other than the 80S ribosome ) also occur in mitochondria and the stroma of chloroplasts. For both of these contexts, relatively large subgroups can be identified in the UMAP. Two further contextual features that may be useful in downstream cryoET analyses are the membrane (a macromolecule segmentation, unlike all other contextual features included in this figure) and void. The former might help differentiate between membrane-bound and unbound ribosomes, whereas the local value of the void segmentations could be used as an exclusion metric (e.g. discarding all particles with a void value above some threshold). For the remaining features (e.g. Golgi, pyrenoid, vesicle, etc.), the number of associated ribosomes was relatively low. That any ribosomes were detected at all in these typically ribosome-free cellular environments is likely due to inaccuracies in the network used for ribosome segmentation. For a purpose such as subtomogram averaging, it may also be useful to discard these particles from the selection.

**Figure 5 supplement 2.**
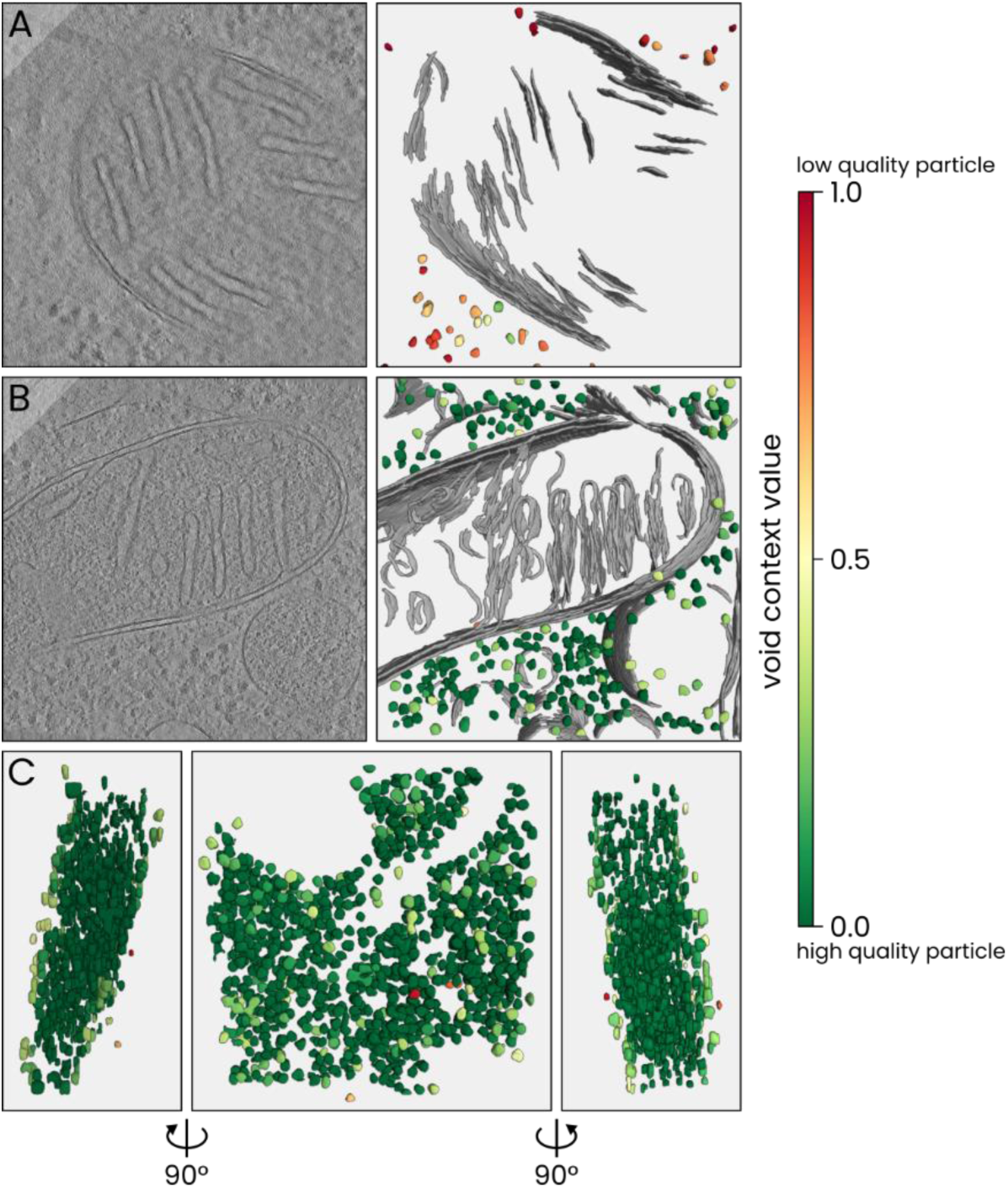
Comparing ribosomes with low and high ‘void’ context values. Ribosome segmentations are rendered as isosurface meshes, with particles coloured in relation to the local void context value. **A)** An example of a relatively low quality tomogram, where a number of ribosomes with a high void score were detected. **B)** A tomogram with similar content (mitochondria and ribosomes), but with a higher quality. Most of the ribosomes detected in this volume scored a lower void context value. **C)** An example to illustrate the spatial distribution of low and high void value ribosomes. The same segmentation is rendered from three points of view. As is typical across the dataset, ribosomes with a high void value are found near the top and bottom surface of the sample. In some cases, particles identified as ribosomes and occuring at these surfaces are in fact ice particles that were falsely segmented as ribosomes.

**Figure 5 supplement 3.**
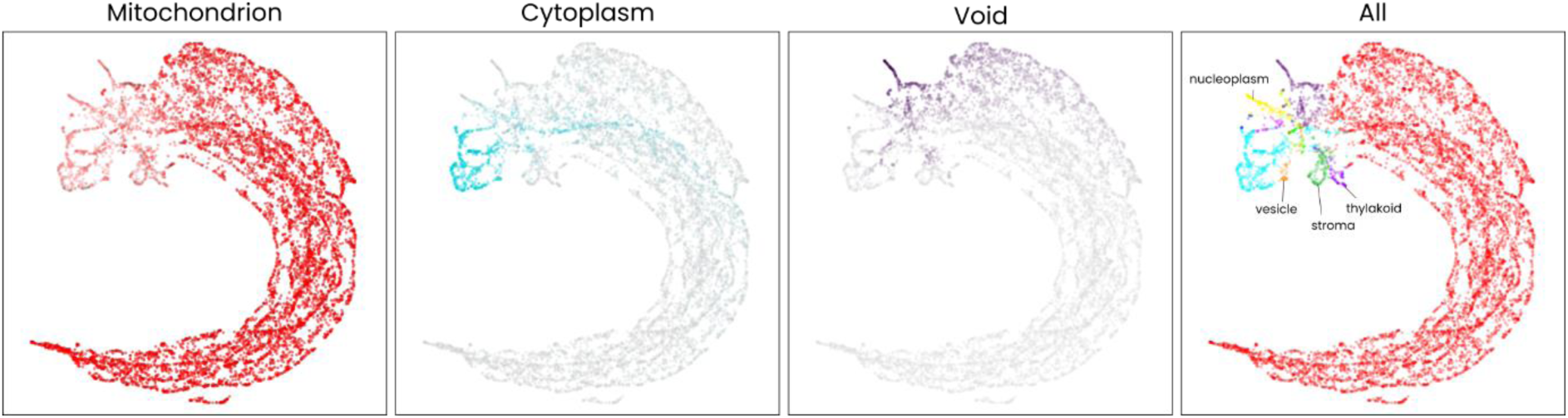
Context-aware particle picking applied to ATP synthase. UMAP of ATP synthase context vectors, coloured by different context values. A total of 8231 ATP synthase molecules were detected across the full dataset (note that ATP synthase is relatively difficult to accurately segment, and that the true number of ATP synthase molecules visible in the data is likely higher). For 6948 (84.4%) of these, the highest context value corresponded to mitochondrion, while 498 scored highest on cytoplasm, 291 on void, and the remaining 494 particles scores highest on any of the remaining features. Since ATP synthase is primarily found within mitochondria (a different type exists within chloroplasts, but was not included in the training dataset for the ATP synthase segmenting neural network and was also rarely recognized in the data, if at all), the first group of 6948 particles is most likely to represent true positive particles, while the remaining 1283 particles are more likely to represent false picks.

**Figure 5 supplement 4.**
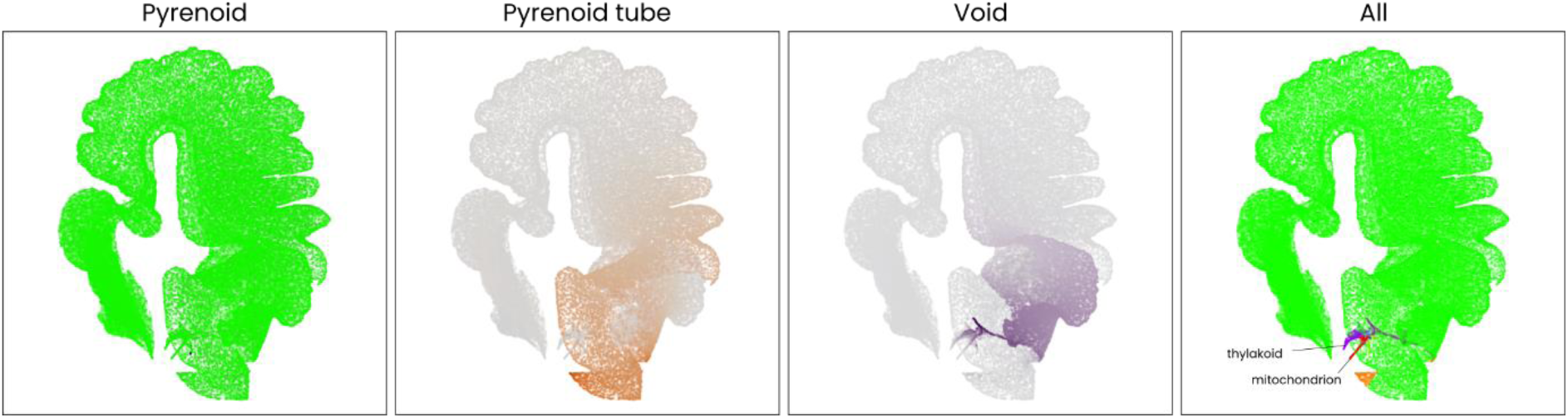
Context-aware particle picking applied to RuBisCo. UMAP of RuBisCo context vectors, coloured by different context values. A total of 244695 RuBisCo molecules were detected across the full dataset. For 241224 (98.6%) of these, the highest context value corresponded to pyrenoid, while a small number scored highest on thylakoid (964, 0.39%), void (838, 0.34%), mitochondrion (595, 0,24%), or pyrenoid tube (353, 0.14%), or any of the other features.

**Figure 6 supplement 1.**
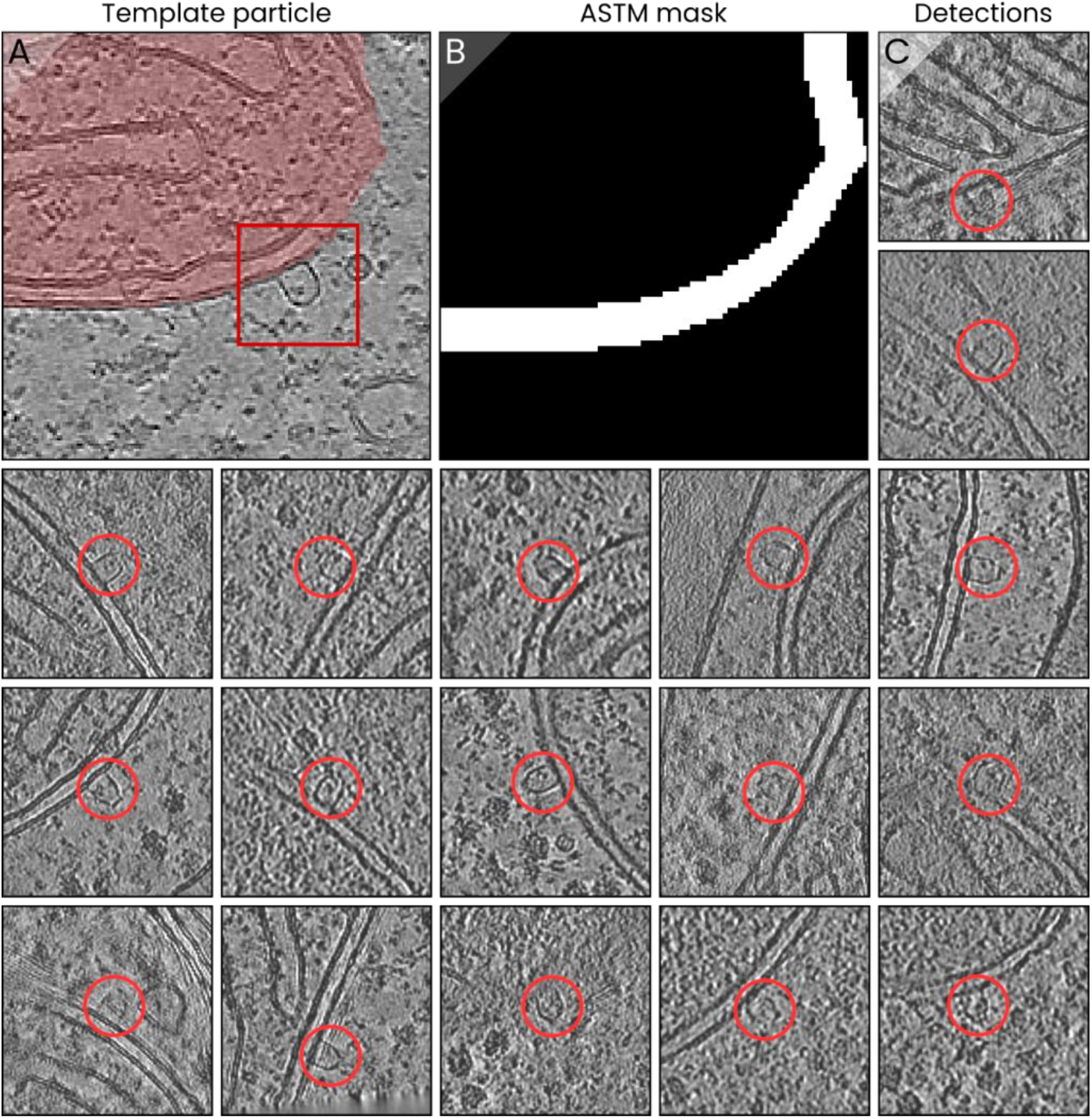
Area-selective template matching applied to ‘lightbulb’ particles. **A)** A single lightbulb particle, which was picked and used as the template. **B)** An example of the volume mask used in ASTM, specifying that TM is to be applied within a ∼20 nm thickness shell outside the mitochondrial outer membrane only. The mask was based on the mitochondrion segmentation. **C)** The resulting lightbulb detections. Since the morphology of the particles appears to vary significantly (in some cases the top – i.e., that furthest away from the membrane – is curved, in other cases distinct edges are visible) and the lumen of these particles resembles the cytoplasm itself, the lightbulbs were relatively difficult targets for detection with TM. The false positive rate in this case was high (∼50 false positives per true particle) and the number of false negatives is likely also high. This problem could be alleviated by improving the implementation of ASTM, or using a better template (using a single particle as the template introduces some difficulty, as the signal to noise ratio is limited). Nevertheless, the resulting set of ∼20 lightbulb particles offers a starting point either for the training of a neural network for lightbulb segmentation, or for generating an initial subtomogram average for use as the template in a subsequent iteration of (AS)TM.

## Supplementary Methods

### Pom python module

The code that was developed during these experiments has been wrapped up into a Python module named Pom, which is now freely available as ‘Pom-cryoET’ on the Python package index. Documentation is available online at pom-cryoet.readthedocs.io.

In combination with Ais, our previously published programme for segmentation of cryoET datasets, **Pom & Ais** enable all processing steps required to perform this workflow. Below we briefly outline the steps involved in the workflow.

The first step is to segment macromolecules. While there are many different tools available to perform this step, we used Ais because we would also use it for annotation in subsequent steps. We refer to the Ais documentation for additional information: ais-cryoet.readthedocs.io

The next step is to train multiple networks that each segment a single organelle. Here we use Ais as an annotation tool. First, we defined each of the chosen organelles in the Ais feature library in order to ensure that the different annotations would remain organised as we went on to annotate each of these organelles in numerous different tomograms.

Next, we opened a number (initially twenty) of tomograms in Ais, browsed through the volumes and between different volumes, and in every instance where we saw an image feature that clearly corresponded to one of the chosen organelles, we would carefully annotate that region and place a box in it. Placing a box means selecting a square region of interest within a tomographic slice; the neural networks receive these square images as the training inputs. While we thus accumulated annotated images for each separate training dataset, we also included boxes that did not contain the image feature of interest (for example: for the pyrenoid training data we would also place a number of boxes in regions of nucleoplasm, cytoplasm, mitochondria, etc.) in order to ensure that the training data supplied to each separate neural network would also include representations of regions that did not represent the feature of interest. After annotating and placing a couple of boxes per a tomogram, we saved the annotated tomogram and continued by annotating another.

Eventually, we had thus prepared a set of sparsely annotated tomograms, which are the input for the next step in the workflow.

The first step when using Pom is to initialize the various training datasets for the single-organelle networks; meaning, to sample all the annotated tomograms as prepared and saved with Ais, and to output files for use in training the various neural networks. This can be done using the following command:

~~~
**pom single initialize**
~~~

Next, the single-organelle networks can be trained, one by one. For example:

~~~
**pom single train -o “Cytoplasm” -gpus 0,1,2,3**
~~~

This typically required ∼30 minutes per network when trained on four GPUs. Using four Linux servers with eight GPUs each, we could train eight networks at the same time, so that the total processing time for preparing all single-organelle networks was typically around 2 – 3 hours.

After training, the following command can be used to test the networks on a subset of tomograms:

~~~
**pom single test -o “Cytoplasm” -gpus 0,1,2,3**
~~~

If the outputs of the single-organelle models are deemed sufficiently accurate (judged by manual inspection or using various validation metrics – we judged by eye), the next step can be started, which is to generate a shared training dataset for training a single network to output all classes simultaneously.

As before, we first initialize the training data:

~~~
**pom shared initialize**
~~~

Next the shared network is trained:

~~~
**pom shared train**
~~~

And after that, the data processed:

~~~
**pom shared process**
~~~

This processing step was implemented with parallel processing on multiple servers in mind. When running the process command simultaneously on different servers, files are dynamically distributed to different workers.

Using the four servers with 8 GPUs each, we could thus process 32 volumes simultaneously. With a typical processing cost per volume per GPU of around one minute, this reduced the average cost per volume to ∼2 seconds. As a result, all volumes could be processed in a little under one hour.

Once all segmentations are completed a number of Pom commands are available for summarizing, rendering, and exploring the data. To measure all volume compositions:

~~~
**pom summarize**
~~~

To generate XY and XZ projection images of all segmentations:

~~~
**pom projections**
~~~

To render segmentations as 3D images, with user-specified compositions*:

~~~
**pom render**
~~~

And finally, to launch the Streamlit appliciation for exploring the results:

~~~
**pom browse**
~~~

Since this streamlit app is browser-based, the results can be easily accessed on any PC that is connected to the same network. To illustrate, the report generated for the dataset used in this article is available online via cryopom.streamlit.app

## Footnotes

* The user can configure multiple named compositions. For example: ‘Macromolecules’, showing membranes, ribosomes, etc.; ‘Mitochondrion’, showing membranes, ATP synthase, and mitochondrion segmentations; ‘Nucleus’, showing membranes, nucleoplasm, and nuclear envelope; etc.

